# Co-expression enrichment analysis at the single-cell level reveals convergent defects in neural progenitor cells and their cell-type transitions in neurodevelopmental disorders

**DOI:** 10.1101/2020.02.13.948315

**Authors:** Kaifang Pang, Li Wang, Wei Wang, Jian Zhou, Chao Cheng, Kihoon Han, Huda Y. Zoghbi, Zhandong Liu

## Abstract

Recent large-scale sequencing studies have identified a great number of genes whose disruptions cause neurodevelopmental disorders (NDDs). However, cell-type-specific functions of NDD genes and their contributions to NDD pathology are unclear. Here, we integrated NDD genetics with single-cell RNA sequencing data to identify cell-type and temporal convergence of genes involved in different NDDs. By assessing the co-expression enrichment pattern of various NDD gene sets, we identified mid-fetal cortical neural progenitor cell development—more specifically, ventricular radial glia-to-intermediate progenitor cell transition at gestational week 10—as a key convergent point in autism spectrum disorder (ASD) and epilepsy. Integrated gene ontology-based analyses further revealed that ASD genes function as upstream regulators to activate neural differentiation and inhibit cell cycle during the transition, whereas epilepsy genes function as downstream effectors in the same processes, offering a potential explanation for the high comorbidity rate of the two disorders. Together, our study provides a framework for investigating the cell-type-specific pathophysiology of NDDs.

## Introduction

Over the past decade, large-scale exome and genome sequencing studies have firmly established that *de novo* protein-altering variants contribute significantly to NDDs, including ASD (Iossifov et al. 2014; De Rubeis et al. 2014; Krumm et al. 2015; Sanders et al. 2015; C Yuen et al. 2017), epilepsy (Allen et al. 2013; EuroEPINOMICS-RES Consortium et al. 2017; Heyne et al. 2018), intellectual disability (ID) (de Ligt et al. 2012; Rauch et al. 2012; Lelieveld et al. 2016), and developmental delay (DD) (Deciphering Developmental Disorders Study 2017). Although hundreds of genes with *de novo* protein-altering mutations in a specific NDD have been identified, each gene accounts only for up to a few cases, demonstrating the high heterogeneity of the underlying genetic landscapes. With the diverse and pleiotropic functions of these disease-associated genes, it is challenging to directly pinpoint disease-specific pathophysiology. However, given the similarity of phenotypic symptoms within each NDD, it is reasonable to hypothesize that disease-causing genes in a specific NDD functionally converge on common brain developmental events. Moreover, NDDs share genetic etiology and comorbidities are frequently found, suggesting that convergences of different NDDs may overlap with each other (Anttila et al. 2018; Lo-Castro and Curatolo 2014). Identification of these convergences will undoubtedly contribute to the mechanistic understanding of NDD pathophysiology and potentially lead to novel treatments.

Several systems-level studies have made significant progress in identifying convergences of NDD genes through integrating NDD genes with functional data, such as gene co-expression and protein-protein interaction (Parikshak et al. 2013; Willsey et al. 2013; Hormozdiari et al. 2015; Chang et al. 2015; Krishnan et al. 2016; Shohat et al. 2017; Lin et al. 2015). For example, Parikshak et al. (2013) applied the weighted gene co-expression network analysis to identify modules of co-expressed genes that are enriched for ASD genes (Parikshak et al. 2013). Their top-down analyses suggest that at the circuit level, ASD genes are enriched in superficial cortical layers and glutamatergic projection neurons during fetal cortical development. Willsey et al. (2013) took a bottom-up approach by focusing on nine high-confidence ASD genes and searching for spatiotemporal conditions in which probable ASD genes co-express with these nine genes (Willsey et al. 2013). Using this strategy, they suggest that glutamatergic projection neurons in deep cortical layers of human mid-fetal prefrontal and primary motor-somatosensory cortex are a key point of ASD gene convergence. Hormozdiari et al. (2015), on the other hand, integrated gene co-expression with protein-protein interaction networks to identify modules that enrich for genes mutated in several NDDs (Hormozdiari et al. 2015). Their results demonstrate that different NDDs share a major point of gene convergence during early embryonic brain development. Although the above mentioned and other studies (Chang et al. 2015; Krishnan et al. 2016; Shohat et al. 2017; Lin et al. 2015) applied different methods, the main conclusions are strikingly similar: a substantial subset of ASD and/or other NDD genes converge in fetal cortical development. In addition, dysfunction of fetal cortical development has also been implicated in other neuropsychiatric disorders including schizophrenia (Gulsuner et al. 2013; Gilman et al. 2012).

The majority of co-expression analyses on NDDs utilized the BrainSpan dataset, a spatiotemporal gene expression data from the developing human brain (Kang et al. 2011). While this dataset is instrumental in assessing transcriptional changes during human brain development, it was collected from bulk brain tissue, making it hard to investigate cell-type-specific co-expression patterns to elucidate the underlying disease mechanisms. Recently, the development of single-cell RNA sequencing (scRNA-seq) technology enabled us to interrogate the transcriptomics at the single-cell level. For instance, Zhong et al. (2018) recently reported the scRNA-seq profiles of more than 2,300 single cells in the developing human prefrontal cortex (Zhong et al. 2018). This kind of data provides an unprecedented opportunity to understand NDD pathophysiology in a cell-type-specific manner.

Here, by integrating disease genes from the four NDDs with the scRNA-seq dataset from the human developing prefrontal cortex, we not only identified disease-specific convergence of NDD genes in specific cell types/stages/transitions but also highlighted the critical cellular processes affected in ASD and epilepsy.

## Results

### Identification of high-confidence genes associated with NDDs

To identify high-confidence risk genes associated with each NDD, we first interrogated genes with *de novo* protein-altering variants for the four NDDs in the denovo-db database (Turner et al. 2017) and non-redundant data for epilepsy (Epi) from two studies (EuroEPINOMICS-RES Consortium et al. 2017; Heyne et al. 2018). Loss-of-function (nonsense, frameshift, and canonical splice site) mutations generally lead to disruption of gene function, whereas missense mutations can cause hypomorphic, hypermorphic, antimorphic, or neomorphic effects. Thus, for each NDD, we divided the associated genes into two categories: genes with *de novo* loss-of-function (dnLoF) mutations and genes with *de novo* missense (dnMis) mutations. To select the most relevant genes for each NDD, we only included genes with at least two or three (depending on gene set sizes) *de novo* mutations of the same category in each specific disorder (see Methods). In total, we defined eight high-confidence NDD gene sets: dnLoF-ASD, dnLoF-Epi, dnLoF-ID, dnLoF-DD, dnMis-ASD, dnMis-Epi, dnMis-ID, and dnMis-DD (**Supplementary Table S1A**). There are some overlaps among different gene sets, which is expected given the high comorbidity among these NDDs (**Supplementary Fig. S1**).

### Different NDD gene sets display distinct co-expression enrichment across major cortical cell types

Previous co-expression analyses on NDDs used transcriptomic data from bulk brain tissue (Parikshak et al. 2013; Willsey et al. 2013; Hormozdiari et al. 2015; Lin et al. 2015). While these analyses are important to identify critical developmental stages and biological processes involved in the specific NDD, it is challenging to dissect cell-type-specific contributions to the disease pathophysiology. Dysfunction of the prefrontal cortex has been implicated in multiple NDDs (Xiong et al. 2007; Gulsuner et al. 2013; Willsey et al. 2013; Arnsten 2006; Parikshak et al. 2013). To investigate the co-expression dynamics of NDD genes in specific cell types during the human prefrontal cortex development, we utilized a recently published scRNA-seq dataset (Zhong et al. 2018) containing more than 2,300 single cells of the developing human prefrontal cortex from gestational weeks (GWs) 8 to 26. Six major cell classes are identified in this dataset: neural progenitor cells (NPCs), excitatory neurons, interneurons, astrocytes, oligodendrocyte progenitor cells (OPCs), and microglia. Thus, we performed co-expression analyses of the different NDD gene sets using the transcriptomic data from each of these cell types.

We reasoned that if mutations in different genes can cause similar symptoms in affected individuals, these genes are more likely to functionally converge at some processes and stages in brain development, potentially within a specific cell type. This functional convergence should be reflected by an increase in the level of co-expression within a particular NDD gene set compared with the overall co-expression level of all the expressed genes (background genes) in that cell type. We first calculated the pairwise Spearman’s correlation coefficients between background genes in each cell type and defined the top 0.5% pairs of genes with the highest correlation coefficients as significant co-expressed gene pairs. We then calculated the fraction of significant co-expressed gene pairs out of all pairs of genes in the NDD gene set and divided it by 0.5% to get a co-expression fold enrichment score of the NDD gene set (see Methods). A high co-expression fold enrichment score of an NDD gene set indicates that the genes in the NDD gene set are more significantly co-expressed than background genes. To verify the enrichment in NDD gene sets is indeed specific and disease-relevant, we also included several control gene sets, including genes with dnLoF mutations in unaffected ASD siblings (Turner et al. 2017), genes with LoF mutations in the general population (Lek et al. 2016), brain-specific gene regulatory factors (Brain-GRF) (Berto et al. 2016), and synaptic genes (Koopmans et al. 2019) (**Supplementary Table S1A**).

We calculated co-expression fold enrichment scores for the eight NDD gene sets and four control gene sets across the six major cell types (**Fig. 1A**; **Supplementary Fig. S2**). In general, NDD gene sets show significantly higher co-expression enrichment than control gene sets (**Fig. 1A**; **Supplementary Fig. S2** and **S3**). Several interesting co-expression enrichment patterns can be found. First, the majority of NDD gene sets show high co-expression enrichment in NPCs, suggesting a convergent involvement of NPCs in different NDDs (**Fig. 1A**). Moreover, dnLoF-ASD and dnMis-Epi genes stand out as having the highest co-expression enrichment scores in particular cell types (**Fig. 1A**; **Supplementary Fig. S4**). Specifically, dnLoF-ASD genes have the highest co-expression in NPCs (18.8-fold enrichment), suggesting a significant contribution of NPCs to ASD pathophysiology (**Fig. 1A**). Interestingly, dnMis-ASD genes show low co-expression enrichment in the six major cell types (**Fig. 1A**). This is consistent with the previous estimation that ∼43% of dnLoF mutations contribute to ASD diagnosis but only ∼13% of dnMis mutations do so (Iossifov et al. 2014). Instead, dnMis-Epi genes are highly co-expressed in NPCs, excitatory neurons, and, more prominently, interneurons (**Fig. 1A**). This is in line with previous findings that dnMis mutations significantly contribute to the etiology of epilepsy (Hamdan et al. 2017; Heyne et al. 2018) and dysfunction in interneurons contributes to the pathophysiology of epilepsy (Lado et al. 2013; Noebels 2015). Compared with ASD and epilepsy genes, ID and DD genes do not exhibit comparable enrichment, suggesting less functional convergences of these disease genes. Collectively, our findings reveal that cell-type-specific functional convergences of NDD genes correlate with the underlying genetic architecture of NDDs.

**Figure 1.**
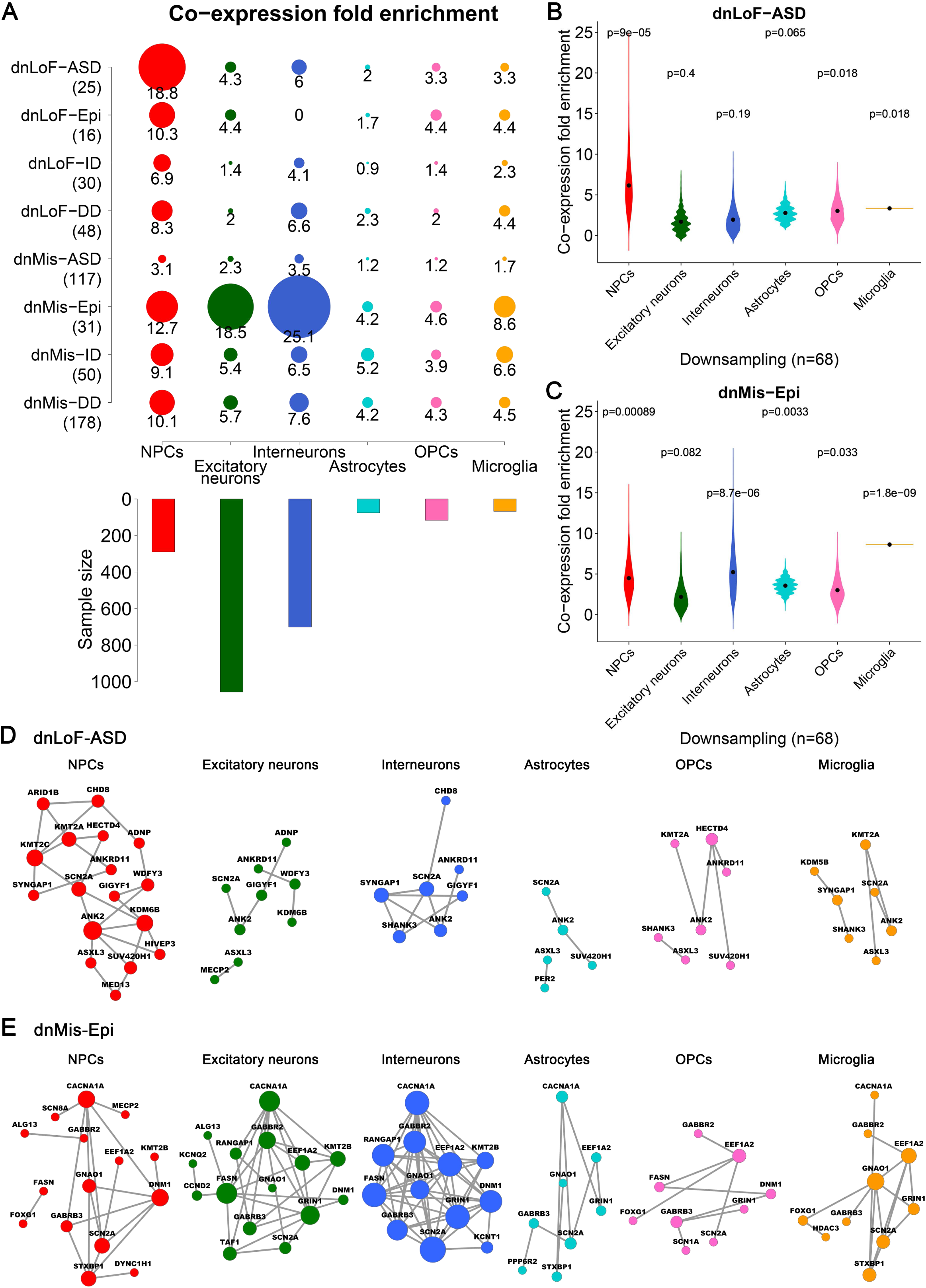
Co-expression enrichment analysis of high-confidence NDD genes in six major cell types of the human prefrontal cortex. (*A*) Co-expression fold enrichment of four NDD gene sets with dnLoF mutations and four NDD gene sets with dnMis mutations in six major cortical cell types as well as the sample size of the cell types. Gene set size is shown in parentheses. Circle size is proportional to co-expression fold enrichment score. (*B,C*) Co-expression fold enrichment of dnLoF-ASD (*B*) and dnMis-Epi genes (*C*) in six major cortical cell types by downsampling the same number of cells for each cell type. The violin plot shows the mean value (point). The statistical significance P value measures whether the mean co-expression fold enrichment score of the corresponding gene set is higher than that of the background genes by the one-sided Fisher’s exact test. (*D,E*) Co-expression networks of dnLoF-ASD (*D*) and dnMis-Epi genes (*E*) in six major cortical cell types using the original sample size. Node size is proportional to co-expression degree.

To determine whether the observed co-expression enrichment reflects true biological signal or is confounded by other factors (Crow et al. 2016; McCall et al. 2016; Skinnider et al. 2019), we systematically tested the possible confounders. We found that the co-expression enrichment is robust to changes in the co-expression threshold (**Supplementary Fig. S5** and **S6**) and correlation-based measures of association (**Supplementary Fig. S7**). The co-expression enrichment also remains similar after controlling for gene set size difference (**Supplementary Fig. S8**), gene expression level dependence (**Supplementary Fig. S9**), and severity of missense mutations (**Supplementary Fig. S10**). Because cell numbers vary across the six major cell types (**Fig. 1A**; **Supplementary Table S1B**), we downsampled the same number of cells for each major cell type to make the co-expression enrichment scores comparable. We found that reducing cell numbers generally decreases the co-expression enrichment scores (**Fig. 1B,C**; **Supplementary Fig. S11**), consistent with the previous finding that larger cell numbers facilitate the reconstruction of more robust and coherent networks (Skinnider et al. 2019). However, even after downsampling, dnLoF-ASD genes still have the highest co-expression in NPCs (**Fig. 1B**), and dnMis-Epi genes are still highly co-expressed in NPCs and interneurons (**Fig. 1C**). Although we used percentile-based cutoff for co-expression enrichment analysis to mitigate the effect of global co-expression differences across cell types, the findings are consistent with results from absolute correlation analysis (**Supplementary Fig. S12** and **S13)**. An unexpected finding is that dnMis-Epi genes have the highest co-expression in microglia after downsampling (**Fig. 1C**). Although microglia have been implicated in epilepsy (Vezzani et al. 2011, 2013), we focused on NPCs and interneurons for further analysis as they have larger sample sizes thus more robust signals.

**Supplementary Fig. S14** and **S15** present several examples of dnLoF-ASD and dnMis-Epi gene pairs that show higher co-expression in NPCs and interneurons, respectively. **Fig. 1D,E** show the co-expression networks for dnLoF-ASD and dnMis-Epi genes in the six major cell types using the original sample size, highlighting the larger number of network edges in the cell types with higher co-expression enrichment.

### ASD and epilepsy genes co-express at specific developmental stages within NPCs and interneurons

Our analyses indicate that the functions of dnLoF-ASD genes converge in NPCs and the functions of dnMis-Epi genes converge in NPCs and interneurons. To determine the specific developmental stages that contribute to the co-expression of dnLoF-ASD and dnMis-Epi genes in NPCs and interneurons, we further performed co-expression enrichment analysis of these two gene sets at different time points. To overcome the effect caused by sample size difference and increase the accuracy of co-expression enrichment score estimation, we focused on cell stages with at least 50 cells and downsampled the same number of cells for each cell stage to make results comparable (**Fig. 2A-C**; **Supplementary Fig. S16**). Apart from NPCs and interneurons where ASD and epilepsy genes show enrichment, we also included excitatory neurons for comparison (**Fig. 2B**).

**Figure 2.**
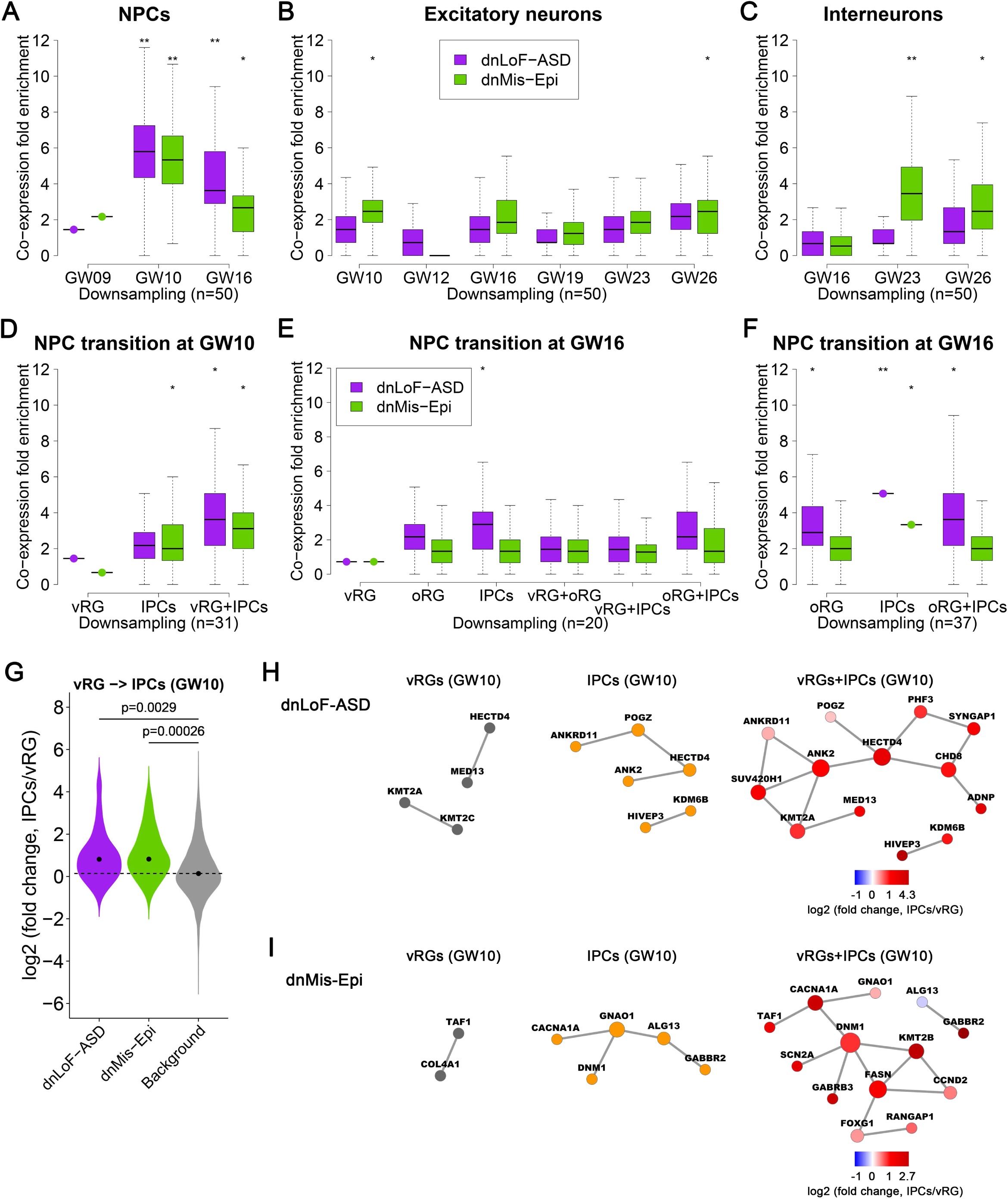
Co-expression enrichment analysis of dnLoF-ASD and dnMis-Epi genes during NPC and neuron development. (*A-C*) Co-expression fold enrichment of dnLoF-ASD and dnMis-Epi genes at specific stages of NPCs (*A*), excitatory neurons (*B*), and interneurons (*C*) by downsampling the same number of cells for each cell stage. (*D*) Co-expression fold enrichment of dnLoF-ASD and dnMis-Epi genes in vRG cells, IPCs, and the transition at GW10 by downsampling the same number of cells for each condition. (*E,F*) Co-expression fold enrichment of dnLoF-ASD and dnMis-Epi genes in vRG cells, oRG cells, IPCs, and their transitions at GW16 by downsampling 20 cells (E) and 37 cells (F) for each condition. In (*A-F*), asterisks above boxplot indicate -log10(P) value that measures statistical significance whether the mean co-expression fold enrichment score of the corresponding gene set is higher than that of the background genes by the one-sided Fisher’s exact test (* 1≤ -log10(P) <2, ** 2≤ -log10(P) <5, *** 5≤ -log10(P) <10, **** 10≤ -log10(P)). (*G*) The expression of dnLoF-ASD and dnMis-Epi genes is significantly increased during the transition from vRG cells to IPCs at GW10. The dashed horizontal line indicates the median log2(fold change) value of the background genes. The statistical significance P values measure whether dnLoF-ASD and dnMis-Epi genes have higher log2(fold change) values than the background genes during the transition by the one-sided Wilcoxon rank sum test. (*H,I*) Co-expression networks of dnLoF-ASD (*H*) and dnMis-Epi genes (*I*) in vRG cells, IPCs, and the transition at GW10 using original sample size. Node size is proportional to co-expression degree.

In NPCs, dnLoF-ASD genes are highly co-expressed at GW10 and, to a lesser extent, GW16 (**Fig. 2A**; **Supplementary Fig. S16A**). GW10 and GW16 are two critical developmental stages for NPCs. NPCs can be further divided into three categories: ventricular radial glia (vRG) cells, outer radial glia (oRG) cells, and intermediate progenitor cells (IPCs) (Lui et al. 2011). The proliferation of IPCs peaks at GW10 and GW16, and they are primarily located in the subventricular zone (SVZ) and outer subventricular zone (oSVZ), respectively (Zhong et al. 2018).

At GW10, vRG cells give rise to IPCs in the SVZ which further differentiate into deep-layer neurons (Nowakowski et al. 2016). Interestingly, dnLoF-ASD genes show little to no co-expression enrichment in vRG cells or IPCs alone at GW10 (**Fig. 2D**; **Supplementary Fig. S17A and S18A**). However, a high co-expression enrichment score was found when vRG cells and IPCs were combined (**Fig. 2D**; **Supplementary Fig. S17A and S18A**). **Supplementary Fig. S19** presents several examples of dnLoF-ASD gene pairs that show high co-expression during the vRG-to-IPC transition at GW10. These results indicate that gene expression variations within vRG cells or IPCs barely contribute to the co-expression of dnLoF-ASD genes in NPCs at GW10. Instead, gene expression variations due to cell-type differences between vRG cells and IPCs largely explain the co-expression enrichment. Consistent with this, we found that the majority of dnLoF-ASD genes concurrently increase their expression during the transition from vRG cells to IPCs at GW10 (**Fig. 2G**; **Supplementary Table S2**). In addition to dnLoF-ASD genes with ≥3 dnLoF mutations, ASD genes with one or two dnLoF mutations and all the SFARI curated gene sets except category six (Basu et al. 2009) also display increased expression during the vRG-to-IPC transition (**Supplementary Fig. S21**). Together, these results highlight the functional convergence of ASD genes in the transition from vRG cells to IPCs at GW10.

At GW16, vRG cells not only give rise to IPCs in the SVZ but also produce oRG cells that will migrate to the oSVZ (Nowakowski et al. 2016; Lui et al. 2011; Fietz et al. 2010; Hansen et al. 2010). In the oSVZ, oRG cells give rise to IPCs that further differentiate into upper-layer neurons (Nowakowski et al. 2016; Lui et al. 2011). We performed similar co-expression enrichment analyses on individual cell types and their combinations. We found that while vRG cells do not show co-expression enrichment, oRG cells and IPCs show moderate co-expression enrichment at GW16 (**Fig. 2E**; **Supplementary Fig. S17B** and **S18B**). However, the co-expression enrichment is not increased in the combination of oRG cells and IPCs, suggesting that gene expression variations both within oRG cells/IPCs and during their transition contribute to the co-expression of dnLoF-ASD genes in NPCs at GW16 (**Fig. 2E,F**; **Supplementary Fig. S17B** and **S18B,C**). Consistently, we found that dnLoF-ASD genes do not show expression change during the transition at GW16 from vRG cells to oRG cells, vRG cells to IPCs, and oRG cells to IPCs (**Supplementary Fig. S22**). Similar results were obtained when analyzing dnMis-Epi genes in NPCs (**Fig. 2A,D-G**; **Supplementary Fig. S16A, S17, S18, S20** and **S22**), whereas the co-expression enrichment score of dnMis-Epi genes at GW16 is generally lower compared with the score of dnLoF-ASD genes at GW16 (**Fig. 2A**). **Figure 2H,I** show co-expression network comparison between individual cell types and the cell-type transition at GW10 for dnLoF-ASD and dnMis-Epi genes using the original sample size.

In excitatory neurons, both dnLoF-ASD and dnMis-Epi genes show moderate to no co-expression enrichment (**Fig. 2B**) despite their elevated absolute correlation at GW16 (**Supplementary Fig. S16B**). In interneurons, dnMis-Epi genes are highly co-expressed at later developmental stages, particularly GW23 (**Fig. 2C**; **Supplementary Fig. S16C**). This coincides with the axon development and cell maturation processes of interneurons in the prefrontal cortex (Zhong et al. 2018).

### Co-expression pattern of ASD and epilepsy genes during the differentiation from NPCs to excitatory neurons

The above analyses focused on co-expression within a major cell type, which mainly captures cell maturation and state changes. To understand whether dnLoF-ASD or dnMis-Epi genes co-function during cell differentiation, we analyzed the co-expression pattern of these two gene sets during NPC terminal differentiation (**Fig. 3A,B**). Due to the sample size limitation (**Supplementary Table S1B**), we focused on the NPC-to-excitatory neuron differentiation at GW10 and GW16 whose time-matched cell stages containing at least 50 samples in both NPCs and excitatory neurons. Excitatory neurons sampled from GW10 and GW16 are mostly deep-layer neurons and upper-layer neurons, respectively (**Supplementary Fig. S23)**. We found that both dnLoF-ASD and dnMis-Epi genes display lower co-expression enrichment in either excitatory neurons or the combination of NPCs and excitatory neurons than in NPCs (**Fig. 3A,B**; **Supplementary Fig. S24**). Also, no co-expression increase was observed during the differentiation from NPC subtypes to excitatory neurons (**Supplementary Fig. S25A,B** and **S26A-C)**. However, both dnLoF-ASD and dnMis-Epi genes tend to increase their expression during the NPC-to-excitatory neuron differentiation, especially at GW16 (**Fig. 3C,D**; **Supplementary Fig. S25C,D** and **S26D-F**; **Supplementary Table S3**). These results suggest that at the individual gene level, ASD and epilepsy genes generally become more abundant/important, yet their functions become more diverse and less convergent in differentiated excitatory neurons than in NPCs.

**Figure 3.**
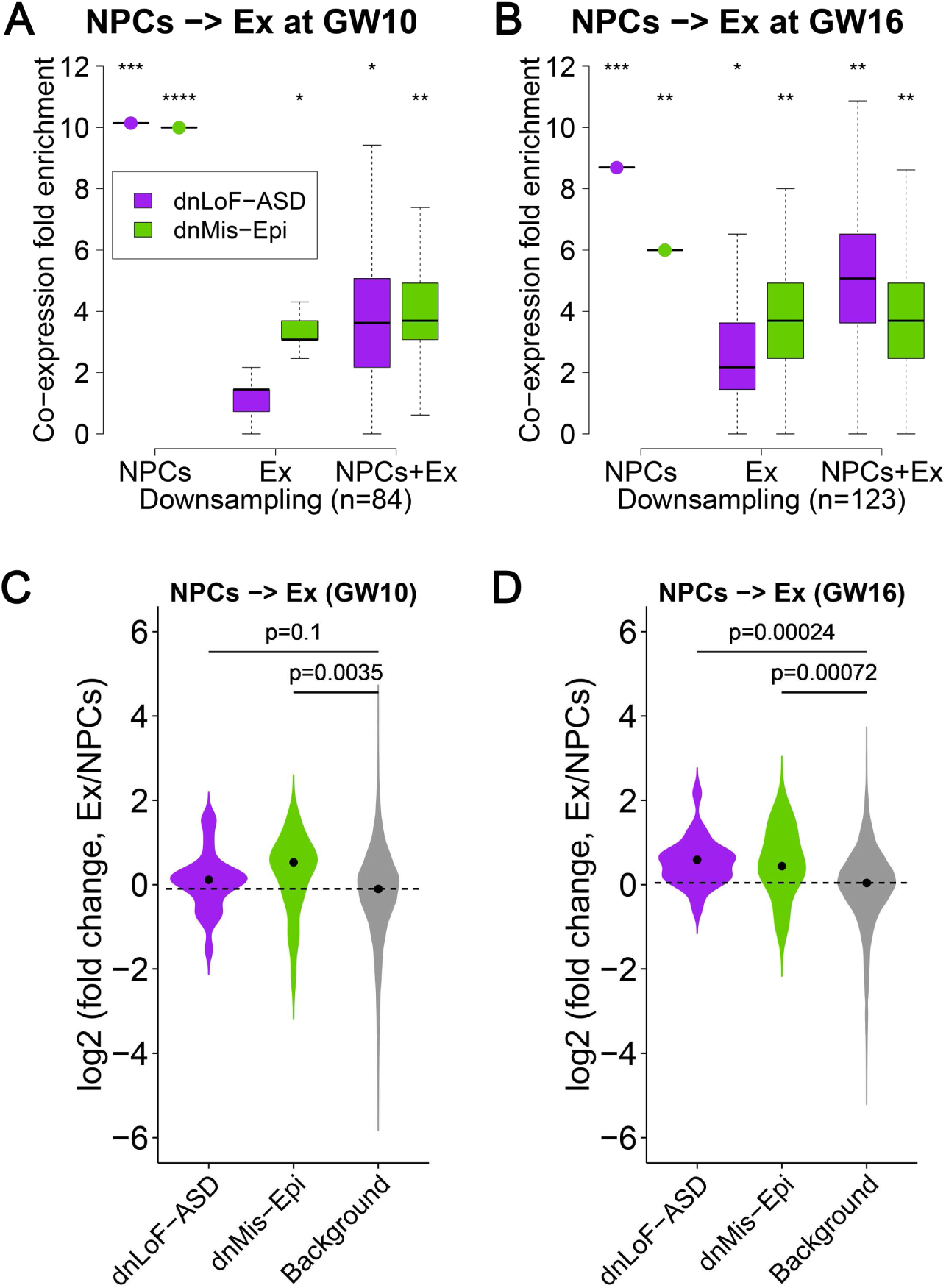
Co-expression enrichment analysis of dnLoF-ASD and dnMis-Epi genes during differentiation from NPCs to excitatory neurons (Ex). (*A,B*) Co-expression fold enrichment of dnLoF-ASD and dnMis-Epi genes in NPCs, excitatory neurons, and the differentiation at GW10 (*A*) and GW16 (*B*) by downsampling the same number of cells for each condition. Asterisks above boxplot indicate -log10(P) value that measures statistical significance whether the mean co-expression fold enrichment score of the corresponding gene set is higher than that of the background genes by the one-sided Fisher’s exact test (* 1≤ -log10(P) <2, ** 2≤ -log10(P) <5, *** 5≤ -log10(P) <10, **** 10≤ -log10(P)). (*C*) The expression of dnMis-Epi but not dnLoF-ASD genes is significantly increased during the differentiation from NPCs to excitatory neurons at GW10. (*D*) The expression of dnLoF-ASD and dnMis-Epi genes is significantly increased during the differentiation from NPCs to excitatory neurons at GW16. In (*C,D*), the dashed horizontal line indicates the median log2(fold change) value of the background genes. The statistical significance P values measure whether dnLoF-ASD and dnMis-Epi genes have higher log2(fold change) values than the background genes during the differentiation by the one-sided Wilcoxon rank sum test.

### Biological pathways associated with ASD and epilepsy genes during the NPC transition at GW10

The above analyses highlight that both dnLoF-ASD and dnMis-Epi genes converge in the vRG-to-IPC transition at GW10. To systematically pinpoint the function of these genes during this transition, we developed a gene ontology (GO) functional analysis method called GO correlation analysis (see Methods). GO correlation analysis was used to determine the correlation between a given gene set and any GO term in a context-dependent manner. Using this method, we calculated Spearman’s correlation for all the GO biological process terms with ASD or epilepsy genes during the vRG-to-IPC transition at GW10. We found that ASD genes are positively correlated with genes involved in neurogenesis and neural differentiation (**Fig. 4A**; **Supplementary Table S4A**) and are negatively correlated with genes involved in cell cycle and cellular respiration (**Fig. 4C**; **Supplementary Table S4C**). Like ASD genes, genes in GO terms that show positive correlation also increase their expression during the transition (**Fig. 4A**; **Supplementary Table S4A**). Instead, genes in GO terms that show negative correlations, especially those involved in the cell cycle, tend to decrease their expression during the transition (**Fig. 4C**; **Supplementary Table S4C**). These observations are consistent with the fact that IPCs exhibit increased neuronal commitment and decreased proliferation capacity compared with vRG cells (Noctor et al. 2004). Similar results were obtained when dnMis-Epi genes were analyzed (**Fig. 4B,D**; **Supplementary Table S4B,D**). These results suggest that both dnLoF-ASD and dnMis-Epi genes are involved in neuron differentiation and cell cycle pathways during the transition.

**Figure 4.**
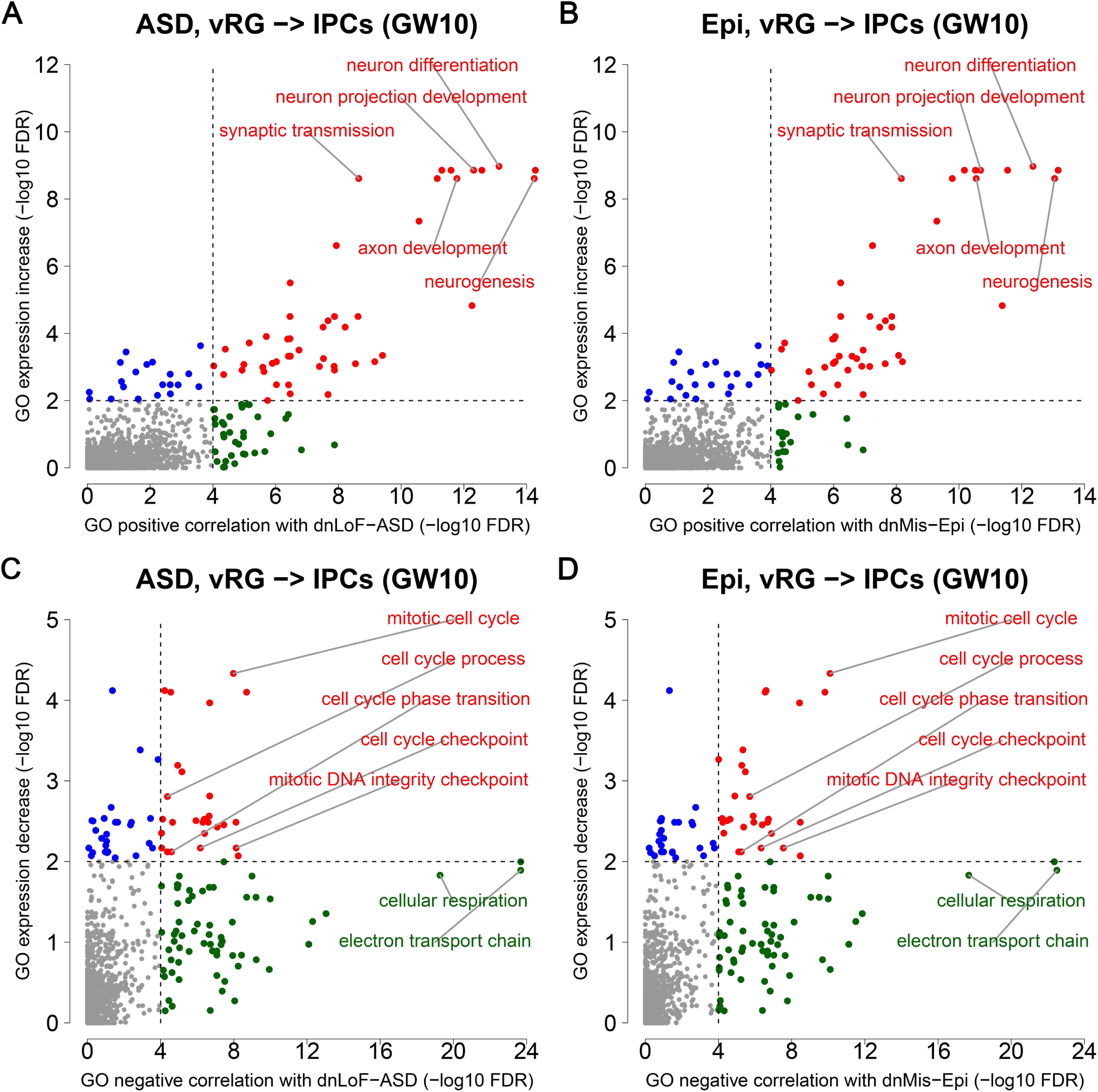
GO correlation and expression change analyses of dnLoF-ASD and dnMis-Epi genes during the vRG-to-IPC transition at GW10. (*A,B*) Scatter plot shows the significance values from GO positive correlation analysis of dnLoF-ASD (*A*) and dnMis-Epi genes (*B*) on the horizontal axis versus the significance values from GO expression increase analysis on the vertical axis during the transition. Dots represent individual GO biological process terms. Each dot has -log10(FDR) value on the horizontal axis that measures how significantly genes annotated under a GO term are positively correlated with dnLoF-ASD (*A*) and dnMis-Epi genes (*B*) during the transition by the one-sided Wilcoxon rank sum test, and - log10(FDR) value on the vertical axis that measures how significantly genes annotated under the GO term have higher log2(fold change) values than the background genes during the transition by the one-sided Wilcoxon rank sum test. The dashed vertical and horizontal lines indicate -log10(FDR) at 4 and 2 as significance thresholds. Significant GO terms from both analyses are shown in red, significant GO terms only from GO positive correlation analysis are shown in green, and significant GO terms only from GO expression increase analysis are shown in blue. Selected representative GO terms are labeled. (*C,D*) Similar to (*A,B*) with GO negative correlation and expression decrease analyses of dnLoF-ASD (*C*) and dnMis-Epi genes (*D*) during the transition.

### Upstream versus downstream involvement of ASD and epilepsy genes during the NPC transition at GW10

It seems that both dnLoF-ASD and dnMis-Epi genes are involved in the same biological pathways during the NPC transition at GW10. However, the manifestations of these two disorders are dissimilar, indicating that the underlying molecular and cellular mechanisms might be different. To determine the difference in ASD versus epilepsy gene functions in NPCs, we examined the composition of each gene set. We found that ASD genes are enriched in GO terms like chromatin modification and organization, but not in the GO terms like neurogenesis and neural differentiation, which are positively correlated with ASD genes (**Fig. 5A**; **Supplementary Table S5A**). Instead, epilepsy genes are both enriched and positively correlated with GO terms like neurogenesis and neural differentiation (**Fig. 5B**; **Supplementary Table S5B**). Given that chromatin modification and organization are critical for transcriptional regulation and dozens of ASD-associated chromatin regulators have well-known regulatory functions in neurogenesis (Ronan et al. 2013; Ernst 2016; Courchesne et al. 2019), these results suggest that ASD genes serve as upstream regulators to control the transcription of other genes in these pathways to promote the NPC transition at GW10. On the other hand, epilepsy genes themselves could be downstream targets regulated by chromatin regulators and thus serve as downstream effectors in the transition. In addition, both ASD and epilepsy genes do not show enrichment with cell cycle-related GO terms that they negatively correlate with (**Fig. 5C,D**; **Supplementary Table S5C,D**). In this respect, ASD genes might also repress the cell cycle through transcriptional regulation.

**Figure 5.**
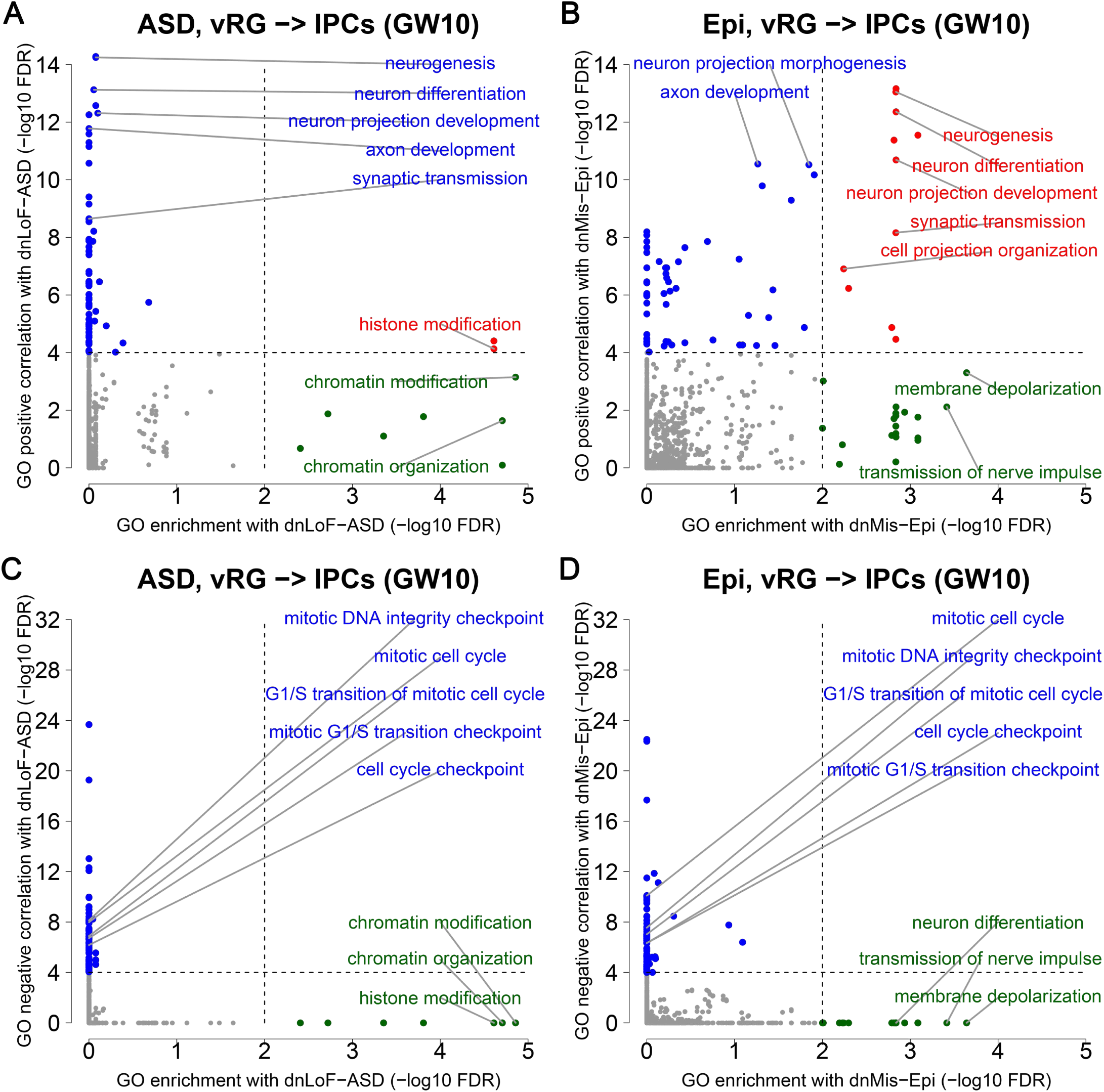
GO enrichment and correlation analyses of dnLoF-ASD and dnMis-Epi genes during the vRG-to-IPC transition at GW10. (*A,B*) Scatter plot shows the significance values from GO enrichment analysis on the horizontal axis versus the significance values from GO positive correlation analysis on the vertical axis of dnLoF-ASD (*A*) and dnMis-Epi genes (*B*) during the transition. Dots represent individual GO biological process terms. Each dot has -log10(FDR) value on the horizontal axis that measures statistical significance of the overlap between genes annotated under a GO term and dnLoF-ASD (*A*) and dnMis-Epi genes (*B*) by the one-sided Fisher’s exact test, and -log10(FDR) value on the vertical axis that measures how significantly genes annotated under the GO term are positively correlated with dnLoF-ASD genes during the transition by the one-sided Wilcoxon rank sum test. The dashed vertical and horizontal lines indicate -log10(FDR) at 2 and 4 as significance thresholds. Significant GO terms from both analyses are shown in red, significant GO terms only from GO enrichment analysis are shown in green, and significant GO terms only from GO positive correlation analysis are shown in blue. Selected representative GO terms are labeled. (*C,D*) Similar to (*A,B*) with GO enrichment and negative correlation analyses of dnLoF-ASD (*C*) and dnMis-Epi genes (*D*) during the transition.

### *CHD8* regulates transcription to promote neural differentiation and inhibit cell cycle

To test if dnLoF-ASD genes are indeed upstream regulators in the NPC transition, we took the chromatin remodeling gene *CHD8*—a key high-confidence ASD gene (Bernier et al. 2014)—as an example. *CHD8* is a hub gene in the vRG-to-IPC transition network at GW10 (**Fig. 2H**). Gompers et al. (2017) generated germline *Chd8* haploinsufficiency mice and performed RNA-seq analysis using forebrain tissue at five developmental stages (E12.5, E14.5, E17.5, P0, and adult) (Gompers et al. 2017). The top 300 downregulated and top 300 upregulated genes in *Chd8* haploinsufficiency versus wild-type mice at each developmental stage were defined as *CHD8*-activated and -repressed genes, respectively (see Methods and **Supplementary Table S6A**). Interestingly, only *CHD8*-activated genes at E14.5 are both preferentially bound by *CHD8* (Gompers et al. 2017) and enriched for ASD genes (**Supplementary Fig. S27**), suggesting that they are more likely genuine *CHD8* target genes that involve in ASD pathology. Thus, we deemed *CHD8*-activated and -repressed genes at E14.5 as *CHD8* target genes in ASD.

We first analyzed the expression pattern of these *CHD8* target genes in human GW10 NPCs. As shown in **Fig. 2H**, *CHD8* doubles its expression during the vRG-to-IPC transition at GW10 (**Supplementary Table S2**). As expected, we observed that *CHD8*-activated target genes also exhibit significant expression increase and *CHD8*-repressed target genes show significant expression decrease compared with the background genes during the transition (**Fig. 6A**; **Supplementary Table S6B**). Consistently, *CHD8* is more positively correlated with *CHD8*-activated target genes and more negatively correlated with *CHD8*-repressed target genes than the background genes (**Fig. 6B**; **Supplementary Table S6C**). Moreover, *CHD8*-activated target genes are enriched with GO terms related to neurogenesis and neuron development (**Fig. 6C**; **Supplementary Table S6D**), whereas *CHD8*-repressed target genes are enriched with GO terms related to cell cycle (**Fig. 6D**; **Supplementary Table S6E**). Together, these results indicate that *CHD8* promotes the vRG-to-IPC transition at GW10 through transcriptionally activating neural differentiation pathways and repressing cell cycle-related processes. Thus, *CHD8* haploinsufficiency could disrupt the vRG-to-IPC transition at GW10 and shift the proliferation-differentiation balance of vRG cells towards proliferation. Indeed, *Chd8* haploinsufficiency mice show an increase in radial glia cells and a decrease in IPCs during embryonic brain development (Gompers et al. 2017).

**Figure 6.**
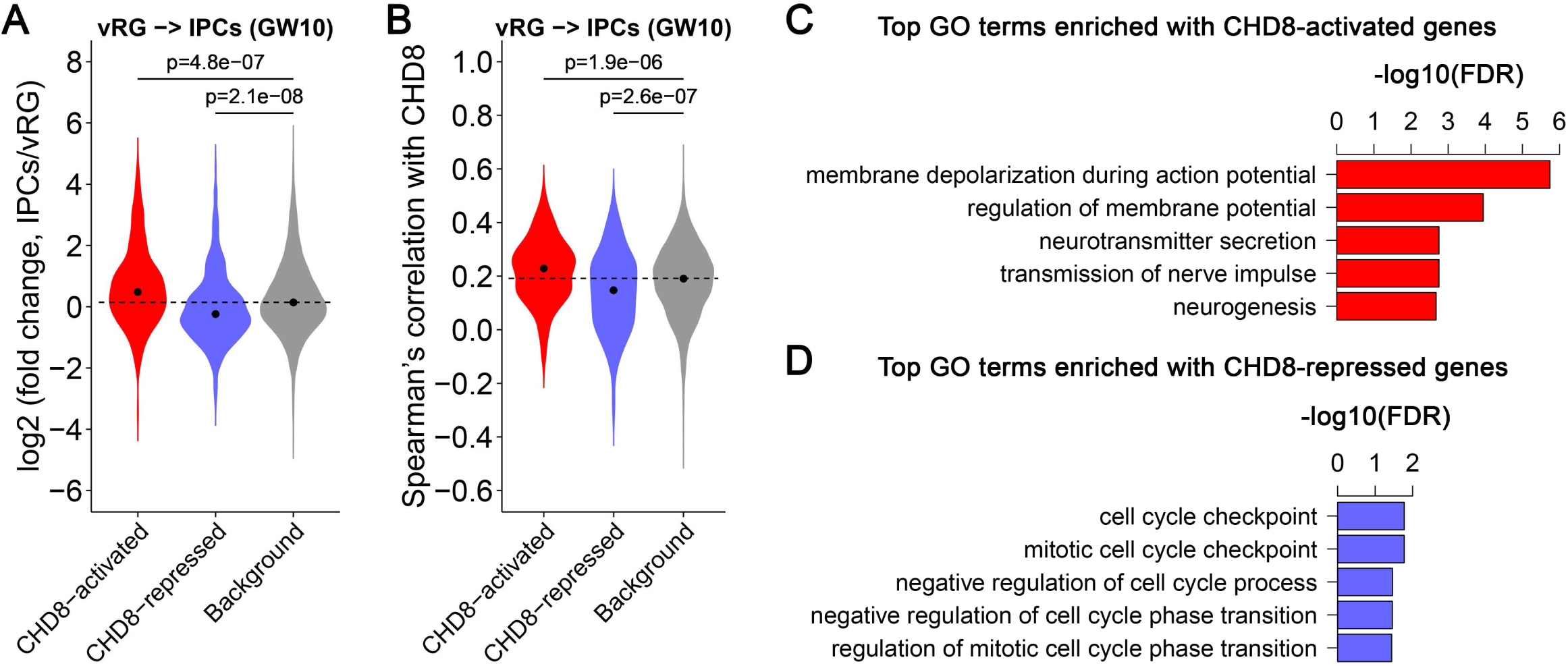
*CHD8* target gene analyses. (*A*) Expression change of *CHD8*-activated and -repressed target genes during the transition from vRG cells to IPCs at GW10. The dashed horizontal line indicates the median log2(fold change) value of the background genes. The statistical significance P values measure whether *CHD8*-activated (-repressed) target genes have higher (lower) log2(fold change) values than the background genes during the transition by the one-sided Wilcoxon rank sum test. (*B*) Spearman’s correlation between *CHD8*-activated/-repressed target genes and *CHD8* during the transition. The dashed horizontal line indicates the median Spearman’s correlation with *CHD8* for the background genes. The statistical significance P values measure whether *CHD8*-activated (-repressed) target genes have higher (lower) correlation with *CHD8* than the background genes during the transition by the one-sided Wilcoxon rank sum test. (*C,D*) Top GO terms enriched with *CHD8*-activated (*C*) and -repressed target genes (*D*).

Collectively, these findings suggest that dnLoF-ASD genes like *CHD8* promote the cell-type transition program by transcriptional regulation of the downstream effectors. On the contrary, dnMis-Epi genes function as effectors that directly participate in the transition processes. Both perturbations would likely affect neural differentiation. However, upstream perturbation by ASD gene mutations could also affect early events of the transition, disrupting the proliferation-differentiation balance of NPCs. Together, these results indicate that mutations of upstream versus downstream genes involved in the same pathways could lead to distinct phenotypic outcomes.

### Co-expression enrichment of NDD genes faithfully represents NDD pathophysiology

All of our co-expression enrichment analyses are based on the assumption that the functional convergences of high-confidence NDD genes represent the core pathways underlying the disease mechanisms. If this assumption is correct, one would expect that low-confidence NDD genes would also converge to the core pathways. To test this possibility, we calculated Spearman’s correlation with dnLoF-ASD genes in NPCs for dnLoF-ASD genes (with ≥3 dnLoF mutations) and ASD genes with fewer dnLoF mutations. As expected, we found that ASD genes harboring two or one dnLoF mutations have a significantly higher correlation with dnLoF-ASD genes than genes harboring no dnLoF mutations, independently validating that the co-expression enrichment of dnLoF-ASD genes in NPCs captures the true ASD pathology (**Fig. 7A**; **Supplementary Table S7A**). Similar results were obtained for dnMis-Epi genes in interneurons (**Fig. 7B**; **Supplementary Table S7B**).

**Figure 7.**
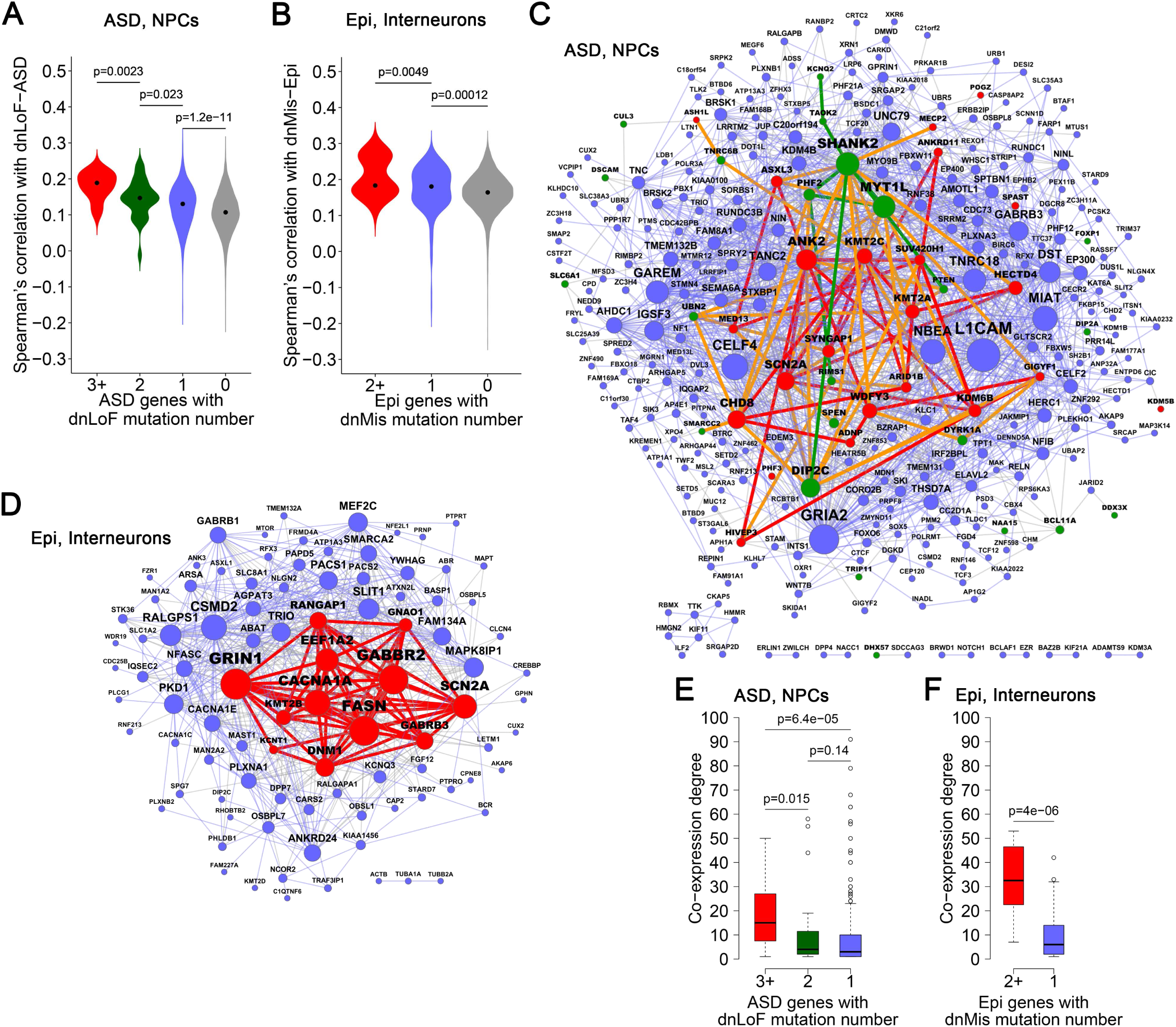
Co-expression network organization of ASD genes with dnLoF mutations in NPCs, and epilepsy genes with dnMis mutations in interneurons. (*A*) Spearman’s correlation with dnLoF-ASD genes in NPCs for ASD genes with ≥3, 2, 1 and 0 dnLoF mutations. (*B*) Spearman’s correlation with dnMis-Epi genes in interneurons for epilepsy genes with ≥2, 1 and 0 dnMis mutations. (*C*) Co-expression network of ASD genes with at least one dnLoF mutations in NPCs. Red, green and blue nodes indicate ASD genes with ≥3, 2 and 1 dnLoF mutations, respectively. Red, green and blue edges indicate co-expression within ASD genes with ≥3, 2 and 1 dnLoF mutations, respectively, and orange edges indicate co-expression between ASD genes with ≥3 dnLoF mutations and ASD genes with 2 dnLoF mutations. (*D*) Co-expression network of epilepsy genes with at least one dnMis mutations in interneurons. Red and blue nodes indicate epilepsy genes with ≥2 and 1 dnMis mutations, respectively. Red and blue edges indicate co-expression within epilepsy genes with ≥2 and 1 dnMis mutations, respectively. In (*C,D*), node size is proportional to co-expression degree. (*E*) Co-expression degree in the NPC network of ASD genes with ≥3, 2 and 1 dnLoF mutations. (*F*) Co-expression degree in the interneuron network of epilepsy genes with ≥2 and 1 dnMis mutations. In (*A,B,E,F*), the statistical significance P values are calculated using the one-sided Wilcoxon rank sum test.

In addition, we found that the Spearman’s correlations with dnLoF-ASD genes in NPCs for dnLoF-ASD genes are significantly higher than those for ASD genes with fewer mutations, and the Spearman’s correlations with dnMis-Epi genes in interneurons for dnMis-Epi genes are significantly higher than those for epilepsy genes with fewer mutations (**Fig. 7A,B**). These results suggest that genes with more mutations tend to be at the core position of the NDD gene co-expression network while genes with fewer mutations tend to be in the peripheral region of the network. To test this hypothesis, we constructed an NPC co-expression network of all the ASD genes with dnLoF mutations (**Fig. 7C**; **Supplementary Table S7C**) and an interneuron co-expression network of all the epilepsy genes with dnMis mutations (**Fig. 7D**; **Supplementary Table S7D**). Consistent with our hypothesis, we found that genes with more mutations tend to be at the core position of the network, as indicated by a significantly higher co-expression degree than genes with fewer mutations (**Fig. 7E,F**; **Supplementary Table S7E,F**). Together, these findings validate that co-expression enrichment of NDD genes faithfully represents NDD mechanisms and provide an explanation of why some NDD genes have more mutations identified than others.

## Discussion

To understand the cell-type-specific mechanisms of NDDs across neurodevelopmental stages, we analyzed the co-expression enrichment patterns of NDD gene sets at the single-cell level. Our results demonstrate that genes that cause different NDDs indeed display distinct co-expression patterns in specific brain cell types. Detailed analyses of subtypes and cell-type transitions at various developmental stages revealed 1) novel convergent functions of dnLoF-ASD and dnMis-Epi genes in the vRG-to-IPC transition at GW10 and 2) novel convergent functions of dnMis-Epi genes in the post-mitotic interneuron maturation. Together, our study supports the hypothesis that heterogeneous genetic mutations in ASD/epilepsy converge to disrupt a small set of critical neurodevelopmental events in particular cell types, expanding our understanding of NDD pathophysiology and stepping towards comprehensive cell maps in neuropsychiatric disorders (Willsey et al. 2018). Our study also presents a computational framework for analyzing disease pathophysiology using scRNA-seq datasets.

### NDD pathophysiology depends on types of genetic perturbations

When analyzing the NDD gene sets, we found that for the same disorder, genes with different types of mutations display distinct co-expression patterns. For instance, dnLoF-ASD genes have the highest co-expression enrichment in NPCs among all the NDD gene sets, but dnMis-ASD genes barely show any enrichment. Instead, dnLoF-Epi genes have the minimum co-expression enrichment in interneurons, while dnMis-Epi genes have the highest enrichment in the same cell type. The exact causes of these observations are not immediately clear. One potential explanation is that haploinsufficiency is the major genetic mechanism for highly penetrant ASD genes. Conversely, gain-of-function or dominant-negative missense mutations dominate the mutational spectrum of highly penetrant genes in epilepsy. Several lines of evidence support this hypothesis: 1) 43% of dnLoF mutations but only 13% of dnMis mutations contribute to ASD diagnosis (Iossifov et al. 2014); 2) dnMis variants explain a larger proportion of individuals with epilepsy than of individuals with ID (Hamdan et al. 2017), and NDD individuals with dnMis variants are more likely to have epilepsy than individuals with dnLoF variants (Heyne et al. 2018); 3) Dozens of dominant-negative or gain-of-function missense mutations have been reported in epilepsy (Yuan et al. 2014; Nava et al. 2014; Orhan et al. 2014; Veeramah et al. 2012; Barcia et al. 2012; Lemke et al. 2014; Li et al. 2016b); 4) At the individual gene level, missense variants in *SCN2A* and *SCN8A* are more strongly implicated in epilepsy than LoF variants (Heyne et al. 2018), and while gain-of-function variants in *SCN2A* contribute to seizure, all ASD-associated variants dampen or eliminate channel function (Ben-Shalom et al. 2017). Nonetheless, whether this hypothesis holds true will require further, more comprehensive investigation.

### NPCs and their cell-type transition in ASD and epilepsy

Another interesting finding is the difference in co-expression patterns within a cell type and during the cell-type transition. We found that both dnLoF-ASD and dnMis-Epi genes are more strongly co-expressed in the whole NPC population than within vRG cells or IPCs alone at GW10. Thus, these genes are less likely to cooperatively function statically in the stemness maintenance or proliferation of vRG cells or IPCs, but convergently play a critical role in the dynamic process of the vRG-to-IPC transition. Consistent with this, most dnLoF-ASD and dnMis-Epi genes, together with other genes critical for neural differentiation, concurrently increase their expression during this transition. Without transcriptomic data at the single-cell level, this kind of subpopulation analysis would be very difficult if not impossible.

The involvement of the vRG-to-IPC transition is interesting. vRG cells are located within the ventricular zone adjacent to the ventricles (Kriegstein and Alvarez-Buylla 2009). vRG cells undergo either symmetric division to proliferate and expand the radial glia pool or asymmetric division to generate neurons or IPCs. IPCs migrate out of the ventricular zone to form the SVZ at the basal side. There, they undergo limited rounds of divisions to produce multiple neurons. It is suggested that this two-step pattern of neurogenesis plays a critical role in the amplification of cell numbers underlying cerebral cortex expansion (Martínez-Cerdeño et al. 2006; Kriegstein et al. 2006). In addition, a perturbation in radial glia cells or IPCs results in abnormal neuron production and cortical malfunction (Krogan et al. 2016; Gompers et al. 2017; Li et al. 2016a; Shenhav et al. 2012). Beyond that, IPCs play an important role in neuronal subtype specification. IPCs, dependent on the time when they are produced, acquire specific neuronal subtype identify and differentially generate cortical layers in a timely manner (Daza et al. 2016). Moreover, the morphological and electrophysiological properties of upper-layer neurons are dependent on their origins from radial glia cells or IPCs (Haydar et al. 2015). Thus, the transition from vRG cells to IPCs has a strong impact on the specificity and function of both the IPCs and the neuronal progeny to be generated.

We found that both ASD and epilepsy genes have higher co-expression enrichment in NPCs than in excitatory neurons. However, their expression levels are higher in excitatory neurons than in NPCs. These findings indicate that at the individual gene level, ASD and epilepsy genes generally become more abundant and potentially function more importantly in young excitatory neurons. However, their functions become more diverse and less convergent in young excitatory neurons as demonstrated by a reduction in co-expression enrichment. Thus, NPCs are likely a more critical convergent point for ASD and epilepsy compared with young excitatory neurons, which could be missed by expression-based analysis (Satterstrom et al. 2020).

### Similar but different roles of ASD versus epilepsy genes during the NPC transition at GW10

We found that ASD genes regulate the transcription of other genes in neural differentiation pathways to promote the NPC transition at GW10. On the other hand, epilepsy genes themselves are downstream effectors controlled by upstream regulators. A mutation in a single ion channel downstream of the differentiation program might severely affect one electrophysiological property of the IPCs, but a mutation in a transcription regulator upstream of the differentiation program could broadly and moderately affect multiple aspects of the cell, such as proliferation, specification, and maturation. Some ASD genes, like *CHD8*, might also determine whether to initiate the transition and/or regulate the balance of NPC proliferation and differentiation at the early stage of the transition. LoF mutations in this kind of genes would promote NPC proliferation at the expense of neural differentiation and cause early brain overgrowth in ASD (Courchesne et al. 2007, 2019; Ernst 2016; Gompers et al. 2017). Some upstream regulators may not only regulate the transition but also specifically control downstream processes related to epilepsy. Mutations in these regulators could lead to both ASD and epilepsy, which may be one reason for such a high comorbidity rate between the two disorders (Sundelin et al. 2016; Betancur 2011).

### An omnigenic model for ASD and epilepsy

The genetic landscapes of ASD and epilepsy are complex and far from completely understood (de la Torre-Ubieta et al. 2016; Cross et al. 2015; Vorstman et al. 2017). With the application of next-generation sequencing and SNP arrays, genetic variations that contribute to the etiology of a number of cases have been uncovered. Still, in most cases, the genetic causes remain unclear. Recently, a new inheritance model for complex diseases—omnigenic inheritance has been proposed (Boyle et al. 2017). In this model, it is suggested that several “core” disease-related genes are responsible for the disease phenotype while all other “peripheral” genes contribute to the phenotype by affecting the functions of these core genes. Due to evolutionary pressure, only a limited number of large-effect genetic variations in core genes can be identified and a large fraction of the total genetic contribution to disease comes from peripheral genes that do not play direct roles. A possible approach to identify core genes is to look for *de novo* rare variants with large effect sizes. This model fits well with our observations that potential core genes with multiple *de novo* rare variants in ASD and epilepsy are clustered at the more central position in the co-expression network of relevant cell types while genes with fewer mutations tend to be in the peripheral region. We noted that another kind of core gene which may function equally importantly across cell types/stages/transitions should not be overlooked. Thus, our study not only provides a list of core genes (such as ASD and epilepsy genes with high co-expression degree in **Supplementary Table S7E,F**) and pathways but also identifies the most relevant cell types where these genes and pathways exhibit convergent function. Future investigations focusing on these core genes and their related regulatory pathways in the most relevant cell types and developmental stages would accelerate NDD gene discovery and enable a more comprehensive understanding of NDD pathophysiology. Development of precise therapies targeting these convergent mechanisms would benefit groups of individuals with NDDs (Sztainberg and Zoghbi 2016; Ernst 2016; Sestan and State 2018; Pang et al. 2014).

### Robustness of co-expression enrichment analysis

Our co-expression enrichment analysis is not affected by confounding factors, such as co-expression threshold, correlation-based measures of association, gene set size, gene expression level, and severity of missense mutations. However, we found that sample size correlates with co-expression enrichment score, and larger cell numbers tend to give higher co-expression enrichment score of an NDD gene set. Based on our observation and the previous finding that larger cell numbers facilitate the reconstruction of more robust and coherent networks (Skinnider et al. 2019), we suggest that controlling for sample size difference be established as a standard for co-expression comparison analysis across different conditions. For the previous conclusions based on co-expression comparison analyses across different conditions without controlling for sample size difference (Willsey et al. 2013; Lin et al. 2015), sample sizes vary across conditions and thus evaluation of sample size effect is probably needed. The potential sample size effect also exists when combining different conditions to construct a global co-expression network, because the signal would be dominated by the conditions with larger sample sizes. Although we used percentile-based cutoff for co-expression enrichment analysis to mitigate the effect of global co-expression differences across cell types, the findings are consistent with results from the absolute correlation analysis. The high co-expression enrichment score also reflects the absolute elevation of co-expression level, especially for dnLoF-ASD genes in NPCs (**Supplementary Fig. S12A**), dnMis-Epi genes in interneurons (**Supplementary Fig. S12B**), dnLoF-ASD and dnMis-Epi genes in NPCs at GW10 and GW16 (**Supplementary Fig. S16A**), and dnLoF-ASD and dnMis-Epi genes in the vRG-to-IPC transition at GW10 (**Supplementary Fig. S18A**). Lastly, it is worth noting that the relatively small sample size has limited our analysis to a few cell types and developmental stages. Besides, we are using the scRNA-seq dataset from the mid-fetal stage of the developing human brain and our analyses primarily focus on early mechanisms of NDDs, that is, transcriptional programs and cell-autonomous effects that take place early in brain development. In the future, it could be fruitful to expand our analysis to more cell types and developmental stages at both cell-autonomous and cell-cell interaction levels when larger scRNA-seq datasets which also cover later developmental stages become available.

## Methods

### High-confidence NDD gene sets

We downloaded *de novo* mutation data for four NDDs: ASD, epilepsy, ID, and DD from the denovo-db v.1.5 database release (Turner et al. 2017) (http://denovo-db.gs.washington.edu). For epilepsy, we also added *de novo* mutation data which were not included in the denovo-db v.1.5 database release from two studies (EuroEPINOMICS-RES Consortium et al. 2017; Heyne et al. 2018). We extracted genes with dnLoF (nonsense, frameshift, and canonical splice site) and dnMis mutations from whole-exome or - genome sequencing data for these four NDDs. The number of dnLoF (dnMis) mutations for a gene in a disorder was defined as the number of distinct individuals with the disorder harboring dnLoF (dnMis) mutations in the gene. High-confidence dnLoF (dnMis) genes for ASD, epilepsy, ID, and DD were defined as genes with at least three dnLoF (dnMis) mutations in each disorder. For high-confidence gene sets with gene number less than 20 (dnLoF-Epi, dnLoF-ID, dnMis-Epi, and dnMis-ID), we used genes with at least two *de novo* mutations. For comparison, we used genes with at least one dnLoF mutations in unaffected ASD siblings in the denovo-db database as sibling control. We also used genes with at least one LoF mutations in the ExAC database (Lek et al. 2016) with known neuropsychiatric cohorts removed as general control. We further included Brain-GRF and synapse genes as controls for genes functioning in the brain. The Brain-GRF gene set is a list of gene regulatory factors that are known to function in the human brain from literature curation (Berto et al. 2016). The synapse gene set was obtained from the SynGO knowledge base (Koopmans et al. 2019). The SFARI ASD gene set was obtained from the SFARI Gene database (Basu et al. 2009), and the SFARI ASD genes were grouped into syndromic genes (category S) and genes with different evidence levels (categories 1-6; high confidence-low evidence). In addition, we assessed whether pathogenicity metrics such as CADD score (Kircher et al. 2014) could improve NDD gene sets with dnMis mutations. We focused on ASD and DD genes with a large number of dnMis mutations available and obtained two high-confidence gene sets: ASD gene sets harboring at least two dnMis mutations with CADD score>25, and DD gene sets harboring at least three dnMis mutations with CADD score>25.

### Processing of scRNA-seq data

The human fetal prefrontal cortical scRNA-seq data (Zhong et al. 2018) used in this study were downloaded from the Gene Expression Omnibus under the accession number GSE104276. The transcript counts of each cell were normalized to transcript per million (TPM), where TPM is the transcript count of each gene divided by the total transcript counts of the cell and multiplied by one million. The gene-level TPM expression values were further transformed to log_2_ (*TPM* + 1) values. Based on the sample annotation file, cells were first divided into six major cell types: NPCs, excitatory neurons, interneurons, astrocytes, OPCs, and microglia. For each cell type, genes with expression level >0 in at least 10% of cells for the cell type were defined as genes expressed in the cell type. Samples in each major cell type were further divided into cell stages based on developmental time points, and only the cell stages containing at least 50 samples were used for analysis. Only the time-matched cell stages containing at least 50 samples in both NPCs and excitatory neurons (astrocytes or OPCs) were used to study the differentiation from NPCs to excitatory neurons (astrocytes or OPCs). Samples in NPCs were further divided into three cell subtypes: vRG cells, oRG cells, and IPCs according to the clustering result of NPCs (Zhong et al. 2018), where vRG cells correspond to clusters 1, 2 and 6, oRG cells correspond to clusters 7, 8 and 9, and IPCs correspond to clusters 3, 4 and 5. Samples in excitatory neurons at GW16 were also divided into three cell subclusters: Ex_C3, Ex_C4, and Ex_C5 according to the clustering result of excitatory neurons (Zhong et al. 2018). The statistical significance P values that measure the expression difference of layer marker genes between GW16 excitatory neuron subclusters and GW10 excitatory neurons were computed using DESeq2 on un-normalized counts (Love et al. 2014). The statistical significance P values of the overlap between eight NDD gene sets were calculated by the one-sided Fisher’s exact test using genes expressed in at least one major cell type as background genes.

### Construction of co-expression networks

To construct a co-expression network for each of six major cell types, we used genes expressed in the cell type as background genes. We first computed the pairwise Spearman’s rank correlation coefficients between background genes and sorted all the pairwise Spearman’s correlation coefficients in descending order. Then, we determined the correlation threshold that gives us the top 0.5% highest pairwise Spearman’s correlation coefficients. The threshold of top 0.5% is commonly used to construct co-expression networks (Lee et al. 2004; Crow et al. 2016) and the value 0.5% was defined as co-expression network density for the background genes. Next, we used the same correlation threshold to construct a co-expression network for a given gene set. For cell stages divided based on developmental time points in each major cell type, we used genes expressed in the major cell type as background genes. For three cell subtypes of NPCs: vRG cells, oRG cells, and IPCs as well as their transitions, we used genes expressed in NPCs as background genes. Genes expressed in either NPCs or excitatory neurons were defined as genes expressed in the NPC-to-excitatory neuron differentiation and used as background genes for the differentiation. The co-expression degree of a gene in the co-expression network is the number of genes co-expressed with the gene. All the co-expression networks were visualized using Cytoscape (Shannon et al. 2003).

### Co-expression enrichment analysis

When constructing a co-expression network for the background genes in one cell type, the value 0.5% used for selection of correlation threshold was defined as co-expression network density for the background genes. Similarly, the co-expression network density for a gene set was defined as the number of significant co-expressed pairs divided by the number of all pairs between genes in the gene set. Then, the co-expression fold enrichment score for the gene set was defined as the ratio of the co-expression network density for the gene set to the co-expression network density for the background genes. The statistical significance of the co-expression fold enrichment score of the gene set was assessed in two ways. First, we compared the co-expression network density for the gene set against the co-expression network density for the background genes by the one-sided Fisher’s exact test with R function:

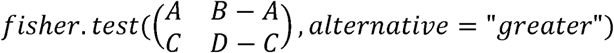

where A is the number of significant co-expressed pairs between genes in the gene set, B is the number of all pairs between genes in the gene set, C is the number of significant co-expressed pairs between the background genes, and D is the number of all pairs between the background genes. Second, we also assessed the statistical significance of the co-expression fold enrichment score of the gene set by comparing whether the gene set has a higher co-expression fold enrichment score than the other NDD gene sets. Similarly, the one-sided Fisher’s exact test was used to compute the statistical significance of the comparison of the co-expression network density for the gene set against the co-expression network density for another NDD gene set.

### Co-expression enrichment analysis by downsampling

Six major cell types have different sample sizes, and microglia has the minimum sample size (68 cells). For fair comparison across the major cell types, we downsampled the same number of cells (68 cells) 1000 times for NPCs, excitatory neurons, interneurons, astrocytes, and OPCs to calculate co-expression fold enrichment score. For fair comparison across the cell stages of the major cell types, we downsampled the same number of cells (50 cells) 1000 times for each cell stage to calculate a co-expression fold enrichment score. During the cell-type transition or differentiation between one cell type with a smaller cell number and the other cell type with a larger cell number, we downsampled the smaller number of cells 1000 times for the cell type with a larger cell number to calculate co-expression fold enrichment score. For the combined cell types, we downsampled half of the smaller number of cells for the cell type with a smaller cell number and half of the smaller number of cells for the cell type with a larger cell number. We then combined the two downsampled cell types and repeated 1000 times to calculate the co-expression fold enrichment score for the combined cell types. To calculate the distribution of average Spearman’s correlation coefficients of an NDD gene set for each condition by downsampling, the pairwise Spearman’s rank correlation coefficients within an NDD gene set were averaged and repeated 1000 times.

### Co-expression enrichment analysis by controlling for different factors

In addition to using the threshold of top 0.5% to construct co-expression networks and calculate co-expression fold enrichment score for NDD gene sets in six major cell types, we used different thresholds of top 0.25% and top 1%. We also varied the thresholds between top 0.1% and top 5% to construct co-expression networks and calculate co-expression fold enrichment score for dnLoF-ASD genes in NPCs and dnMis-Epi genes in interneurons. In addition to using Spearman’s correlation to construct co-expression networks and calculate co-expression fold enrichment score at the threshold of top 0.5% for dnLoF-ASD genes in NPCs and dnMis-Epi genes in interneurons, we used another 16 measures of association implemented in the ‘dismay’ R package (Skinnider et al. 2019). Moreover, we assessed the effect of gene set size difference on the co-expression fold enrichment score of NDD and control gene sets in six major cell types. For each major cell types, we first determined the smallest gene set size of NDD and control gene sets with genes expressed in the cell type. We then downsampled the same number of genes (the smallest gene set size) 1000 times for each gene set to calculate the co-expression fold enrichment score. We further evaluated the dependence of gene expression on the co-expression fold enrichment score of NDD gene sets in six major cell types. For each major cell type, genes expressed in the cell type were divided into ten bins based on expression level with each bin containing the equal number of genes. For each gene set in each cell type, the co-expression enrichment score was computed using 1000 randomly chosen same-size gene sets with the same expression distribution across bins in the cell type as the background gene set.

### Correlation with dnLoF-ASD and dnMis-Epi genes

For the calculation of correlation with dnLoF-ASD genes in NPCs, we used genes expressed in NPCs as background genes. For any non-dnLoF-ASD gene expressed in NPCs, the correlation with dnLoF-ASD genes for the gene was defined as the average Spearman’s correlation coefficients between the gene and dnLoF-ASD genes. For any dnLoF-ASD gene expressed in NPCs, the correlation with dnLoF-ASD genes for the gene was defined as the average Spearman’s correlation coefficients between the gene and the other dnLoF-ASD genes. Based on the correlation with dnLoF-ASD genes for any gene expressed in NPCs, we then obtained the distribution of correlations with dnLoF-ASD genes for different types of mutated ASD genes. The differences in correlations between different ASD gene sets were estimated using the one-sided Wilcoxon rank sum test. A similar analysis was performed to compute the correlation with dnLoF-ASD genes during the transition from vRG cells to IPCs at GW10 using genes expressed in NPCs as background genes. A similar analysis was performed to compute the correlation with dnMis-Epi genes in interneurons and the transition from vRG cells to IPCs at GW10 using genes expressed in interneurons and NPCs as background genes, respectively.

### GO enrichment analysis of dnLoF-ASD and dnMis-Epi genes

To perform GO enrichment analysis, the ontology and human annotation files were downloaded from the GO database (http://www.geneontology.org). To compute the overlap between dnLoF-ASD genes and GO biological process terms during the transition from vRG cells to IPCs at GW10, we used genes expressed in NPCs as background genes. Genes that are annotated under the GO terms but not expressed in NPCs were removed. Only GO terms with the remaining gene number between 10 and 1000 after filtering were used for GO enrichment analysis. The statistical significance P values of the overlap between dnLoF-ASD genes and GO terms were computed using the one-sided Fisher’s exact test and corrected for multiple hypothesis testing using false discovery rate (FDR) control procedure (Benjamini and Hochberg 1995). For GO enrichment analysis of dnMis-Epi genes, the same process above was repeated.

### GO correlation analysis of dnLoF-ASD and dnMis-Epi genes during the cell-type transition

Based on the correlation with dnLoF-ASD genes during the vRG-to-IPC transition at GW10 for any gene expressed in NPCs, we then obtained the distribution of correlations with dnLoF-ASD genes during the transition for genes annotated under a GO biological process term. Only GO terms with the remaining gene number between 10 and 1000 after filtering by genes expressed in NPCs were used. Then, we computed the statistical significance P value which measures whether genes annotated under the GO term have higher correlations than the background genes (genes expressed in NPCs) by the one-sided Wilcoxon rank sum test. We used this P value to measure how significantly the GO term is positively correlated with dnLoF-ASD genes during the vRG-to-IPC transition. We also computed the statistical significance P value which measures whether genes annotated under the GO term have lower correlations than the background genes (genes expressed in NPCs) by the one-sided Wilcoxon rank sum test. We used this P value to measure how significantly the GO term is negatively correlated with dnLoF-ASD genes in the vRG-to-IPC transition. The P values for all GO terms from GO positive or negative correlation analysis of dnLoF-ASD genes during the transition were adjusted using the Benjamini and Hochberg method. For GO correlation analysis of dnMis-Epi genes during the vRG-to-IPC transition, the same process above was repeated.

### Expression change of dnLoF-ASD and dnMis-Epi genes during cell-type transitions

To compute the log2(fold change) value for a gene during the transition from vRG cells to IPCs at GW10, gene expression TPM values of the gene in the vRG and IPC samples at GW10 were added by 1. Then, the average expression of the gene across samples in IPCs at GW10 was divided by the average expression of the gene across samples in vRG cells at GW10 and then log2 transformed. Based on the log2(fold change) value for any gene, we then obtained the distribution of log2(fold change) values for dnLoF-ASD or dnMis-Epi genes. Next, we computed the statistical significance P value which measures whether dnLoF-ASD or dnMis-Epi genes have higher (expression increase) log2(fold change) values than the background genes (genes expressed in NPCs) during the transition by the one-sided Wilcoxon rank sum test. A similar analysis was performed to compute the statistical significance of expression change for dnLoF-ASD and dnMis-Epi genes during the differentiation at GW10 from NPCs, vRG, and IPCs to excitatory neurons, and during the differentiation at GW16 from NPCs, vRG, oRG, and IPCs to excitatory neurons.

### GO expression change analysis during the cell-type transition

Based on the log2(fold change) value for any gene during the transition from vRG cells to IPCs at GW10, we then obtained the distribution of log2(fold change) values for genes annotated under a GO biological process term. Only GO terms with the remaining gene number between 10 and 1000 after filtering by genes expressed in NPCs were used. Then, we computed the statistical significance P value which measures whether genes annotated under the GO term have higher (expression increase) or lower (expression decrease) log2(fold change) values than the background genes (genes expressed in NPCs) by the one-sided Wilcoxon rank sum test. The P values for all GO terms from GO expression change analysis during the transition were adjusted using the Benjamini and Hochberg method.

### *CHD8* target gene analyses

The analytic results of *Chd8* haploinsufficiency mice RNA-seq data were obtained from Supplementary Table S3 of the study (Gompers et al. 2017). Only genes in the *Chd8* RNA-seq data (with gene *CHD8* removed) that are also expressed in NPCs in the human cortical scRNA-seq data were defined as background genes for *CHD8* target gene analyses. The top 300 downregulated and top 300 upregulated genes based on log2(fold change) values in *Chd8* haploinsufficiency versus wild-type mice at each developmental stage were defined as *CHD8*-activated and -repressed genes, respectively. *CHD8*-bound genes are genes whose promoters are bound by *Chd8* in adult mouse forebrain identified using ChIP-seq (Gompers et al. 2017). To compute the overlap between *CHD8*-activated/-repressed genes and *CHD8*-bound genes, we only used *CHD8*-bound genes that are also in the background gene set. The statistical significance P values of the overlap between *CHD8*-activated/-repressed genes and *CHD8*-bound genes were computed using the one-sided Fisher’s exact test. To compute the overlap between *CHD8*-activated/-repressed genes and ASD genes with at least one dnLoF mutations, we only used ASD genes that are also in the background gene set. The statistical significance P values of the overlap between *CHD8*-activated/-repressed genes and ASD genes were computed using the one-sided Fisher’s exact test. Based on the log2(fold change) value for any gene during the transition from vRG cells to IPCs at GW10, we then obtained the distribution of log2(fold change) values for *CHD8*-activated or -repressed target genes. Then, we computed the statistical significance P values which measure whether *CHD8*-activated (-repressed) target genes have higher (lower) log2(fold change) values than the background genes during the transition by the one-sided Wilcoxon rank sum test. Next, we computed the Spearman’s correlation coefficient between any background gene and *CHD8* during the transition from vRG cells to IPCs at GW10 and obtained the distribution of correlations with *CHD8* for *CHD8*-activated or -repressed target genes. We then computed the statistical significance P values which measure whether *CHD8*-activated (-repressed) target genes have higher (lower) correlations with *CHD8* than the background genes during the transition by the one-sided Wilcoxon rank sum test. To compute the overlap between *CHD8*-activated/-repressed target genes and GO biological process terms, we only used GO terms with the remaining gene number between 10 and 1000 after filtering by the background genes. The statistical significance P values of the overlap between *CHD8*-activated/-repressed target genes and GO terms were computed using the one-sided Fisher’s exact test.

## Code availability

Code used in this study is available as **Supplementary Code**.

## Acknowledgments

We thank Shu Zhang and Fuchou Tang for kindly sharing the detailed clustering result of cell subtypes, which can be downloaded now from the Gene Expression Omnibus under the accession number GSE104276. We thank Mingshan Xue, Dmitry Velmeshev, Hyun-Hwan Jeong, and Ying-Wooi Wan for valuable discussions. This work has been supported by National Institute of General Medical Sciences R01-GM120033, National Science Foundation–Division of Mathematical Sciences DMS-1263932, Cancer Prevention and Research Institute of Texas RP170387, Houston Endowment, the Hamill Foundation, and Chao Family Foundation (Z.L.), Huffington Foundation, Howard Hughes Medical Institute (H.Y.Z.). L.W. was supported by a predoctoral fellowship from Autism Speaks (#9120).

## Author contributions

K.P., L.W., H.Y.Z., and Z.L. conceived of and designed the study. K.P. performed analyses. All the authors interpreted the results. K.P., L.W., H.Y.Z., and Z.L. wrote the manuscript with input from W.W., J.Z., C.C., and K.H..

