## Supplementary_Figures for "Co-expression enrichment analysis at the single-cell level reveals convergent defects in neural progenitor cells and their cell-type transitions in neurodevelopmental disorders"

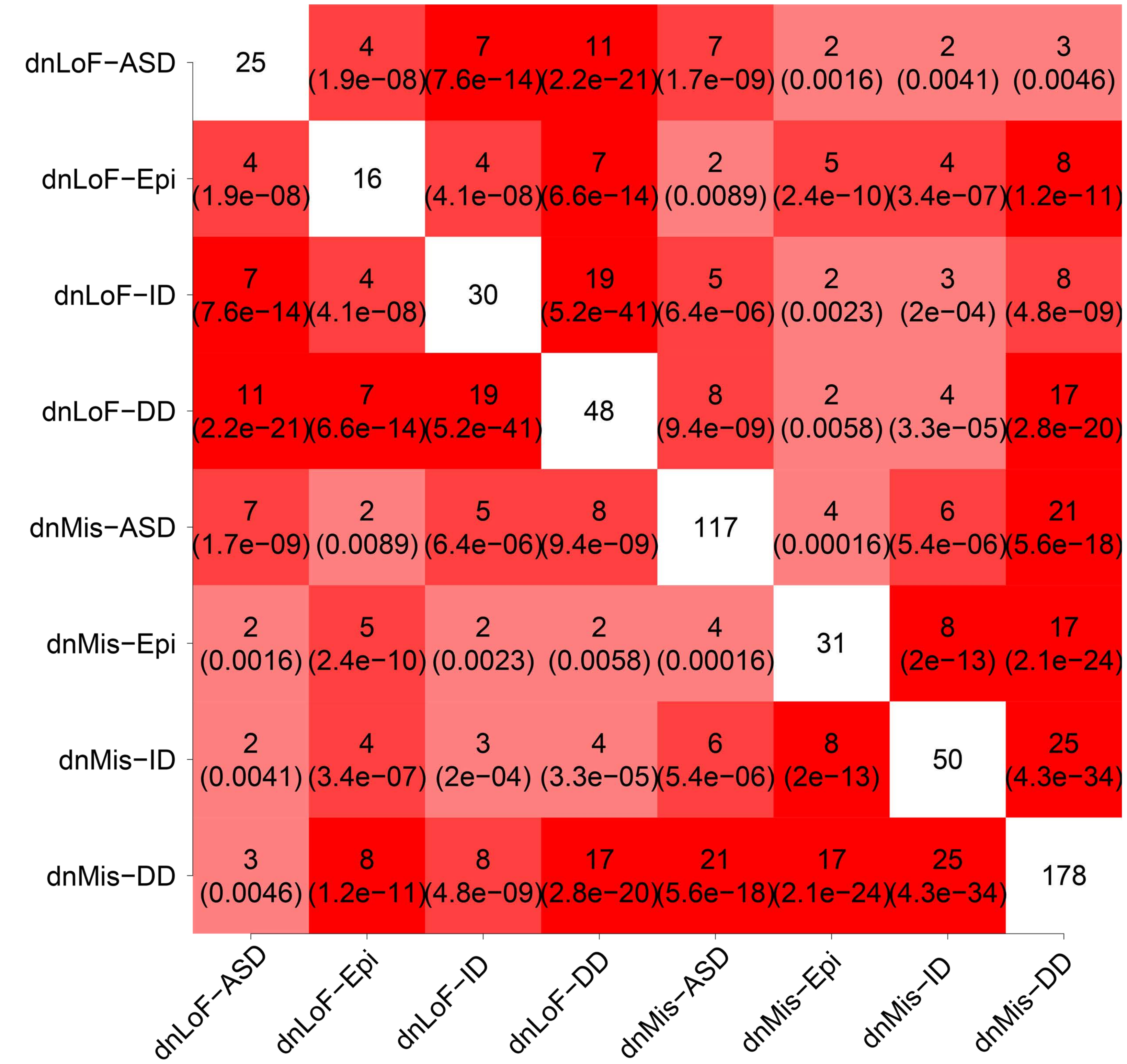


**Supplementary Figure S1.** Overlap between eight NDD gene sets. Square matrix shows the number of overlapping genes between two NDD gene sets and the associated statistical significance P value calculated by the one-sided Fisher's exact test.

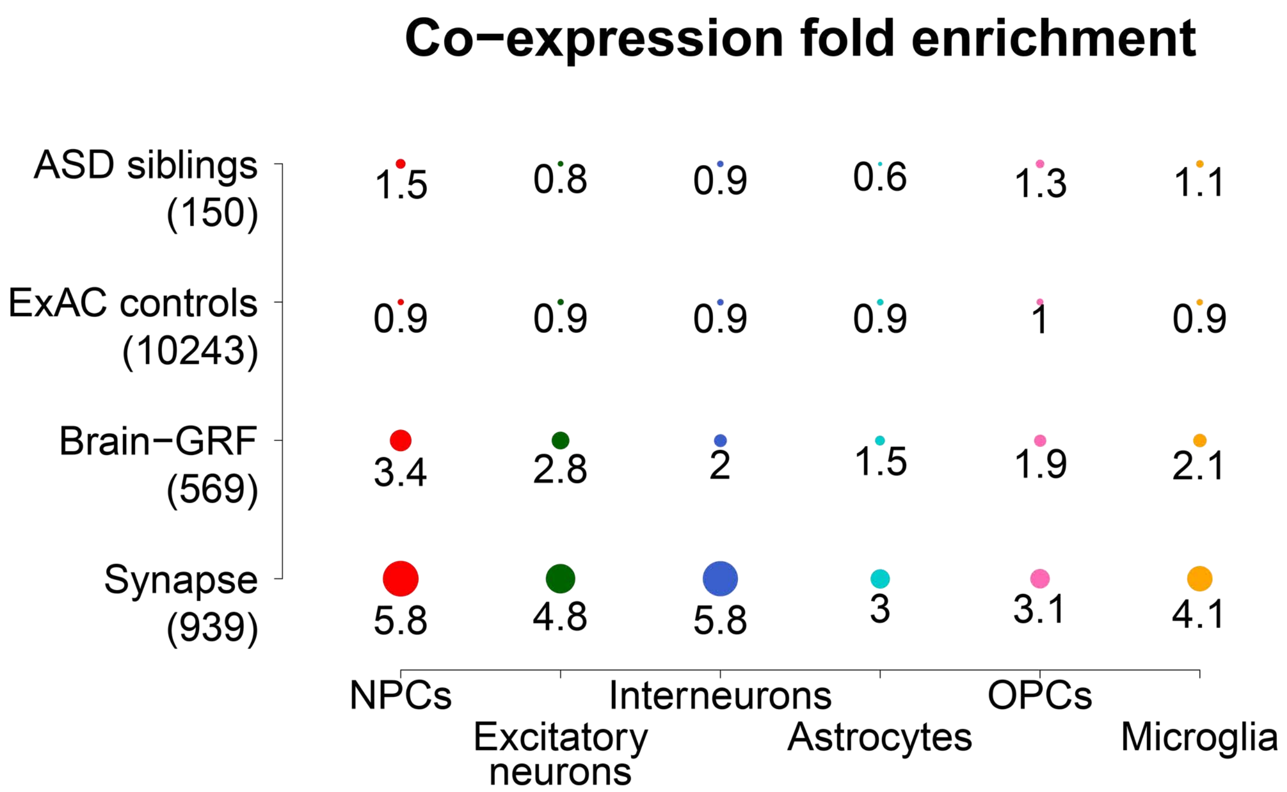


**Supplementary Figure S2.** Control gene sets show low co-expression enrichment in six major cell types of human prefrontal cortex. Co-expression fold enrichment of four control gene sets in six major cortical cell types. Gene set size is shown in parentheses. Circle size is proportional to co-expression fold enrichment score.

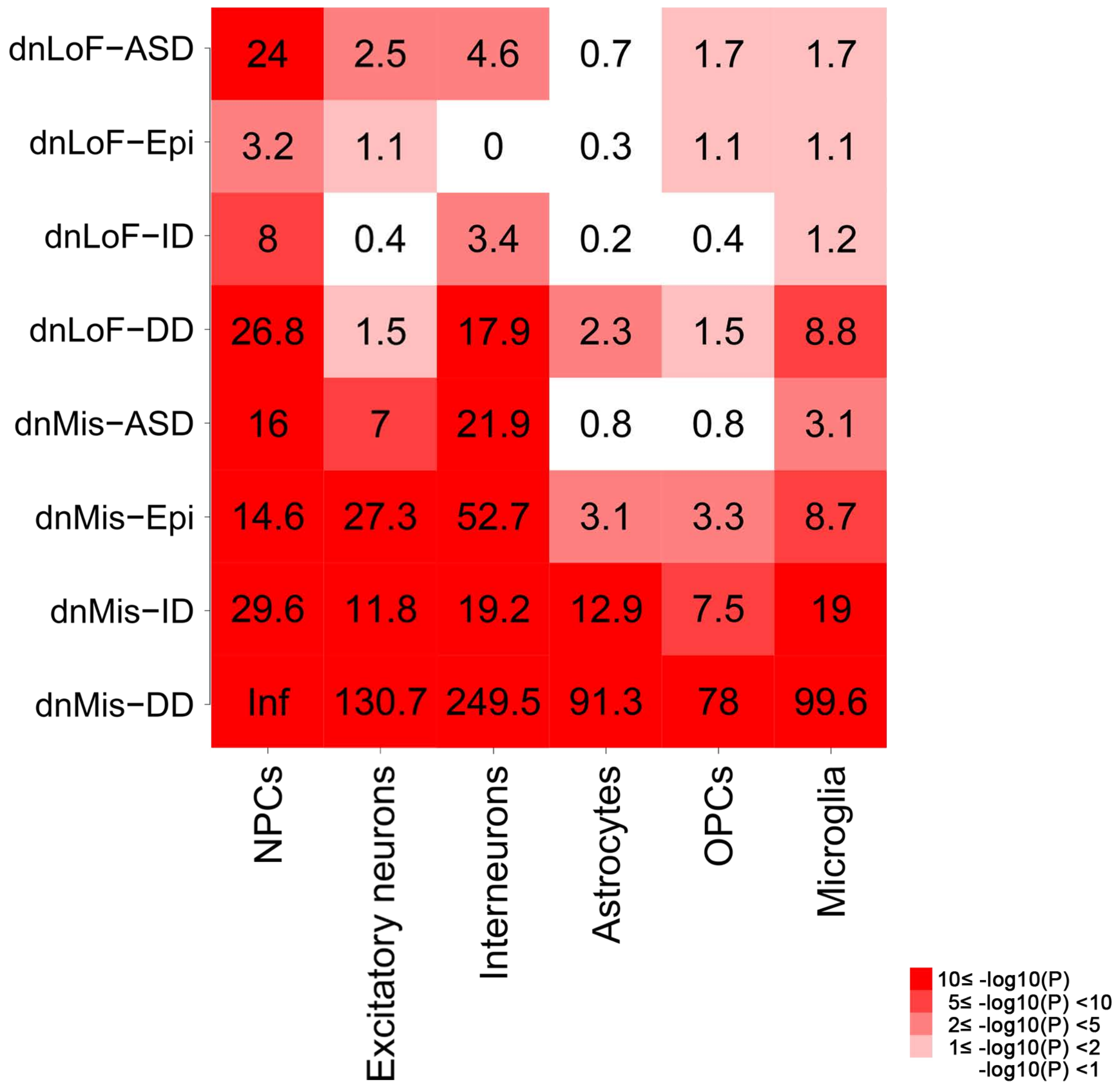


**Supplementary Figure S3.** Significance values of co-expression enrichment analysis of high-confidence NDD genes compared with the background genes. The numerical value in rectangular matrix indicates -log10(P) that measures statistical significance whether the NDD gene set in the corresponding row has a higher co-expression fold enrichment score than the background genes of the cell type in the corresponding column by the one-sided Fisher's exact test.

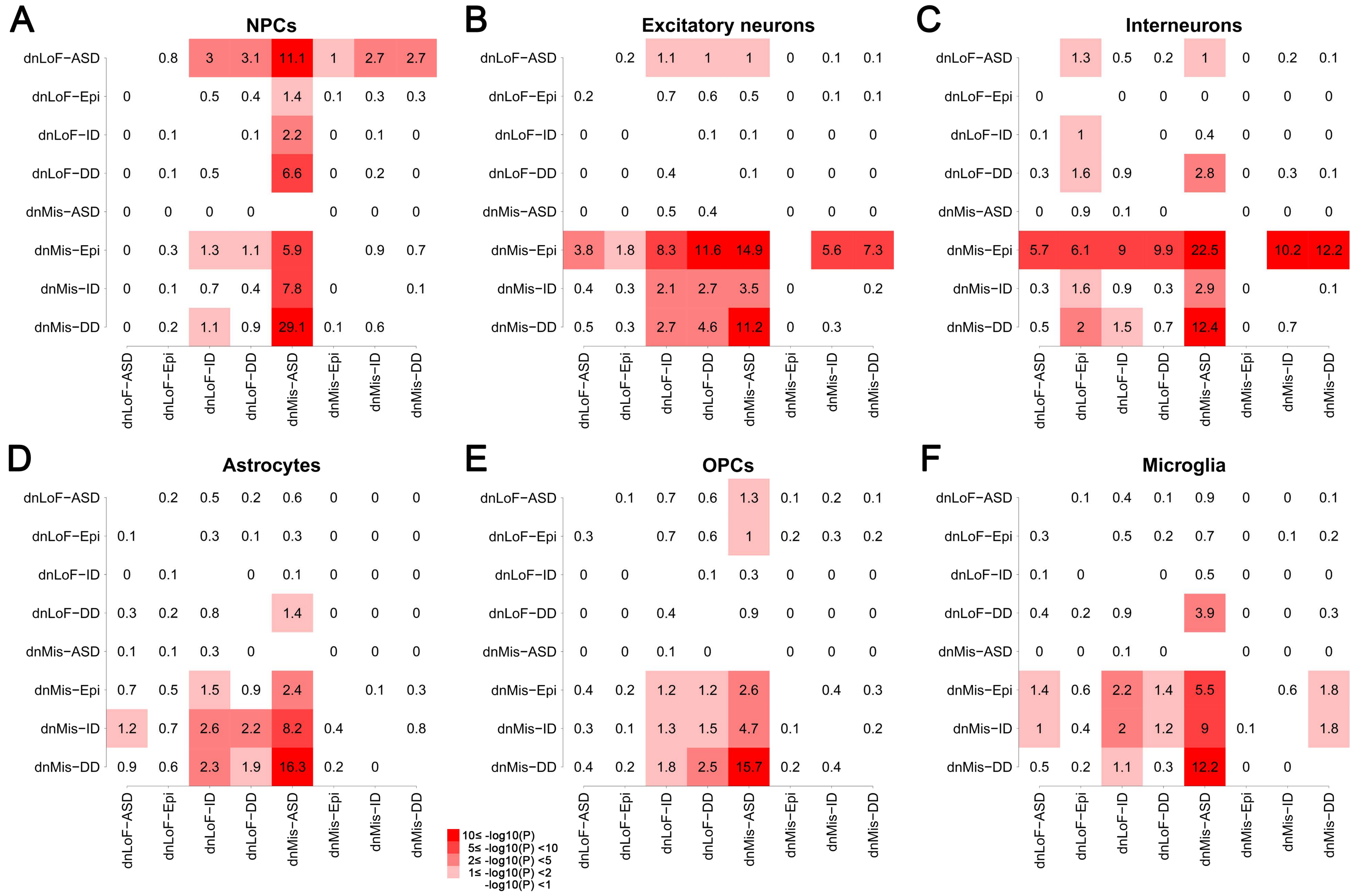


**Supplementary Figure S4.** Significance values of pairwise co-expression fold enrichment comparison between eight NDD gene sets. (*A-F*) Square matrix shows the significance values of the pairwise co-expression fold enrichment comparison between eight NDD gene sets in NPCs (*A*), excitatory neurons (*B*), interneurons (*C*), astrocytes (*D*), OPCs (*E*), and microglia (*F*). The numerical value in each square matrix indicates -log10(P) that measures statistical significance whether the NDD gene set in the corresponding row has a higher co-expression fold enrichment score than the NDD gene set in the corresponding column by the one-sided Fisher's exact test.

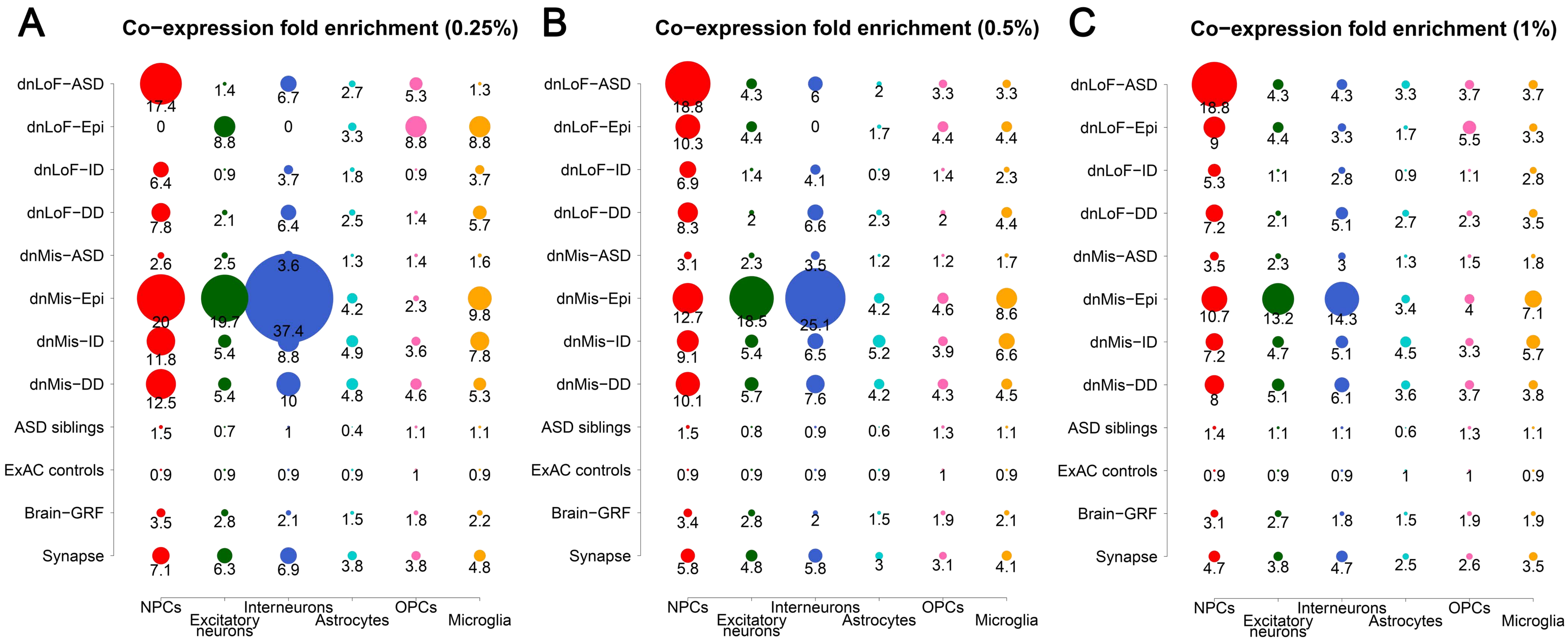


**Supplementary Figure S5.** Co-expression enrichment analysis of twelve gene sets in six major cell types of human prefrontal cortex at different co-expression thresholds. (*A-C*) Co-expression fold enrichment of eight NDD gene sets and four control gene sets in six major cortical cell types at co-expression thresholds of top 0.25% (*A*), top 0.5% (*B*), and top 1% (*C*). Circle size is proportional to co-expression fold enrichment score.

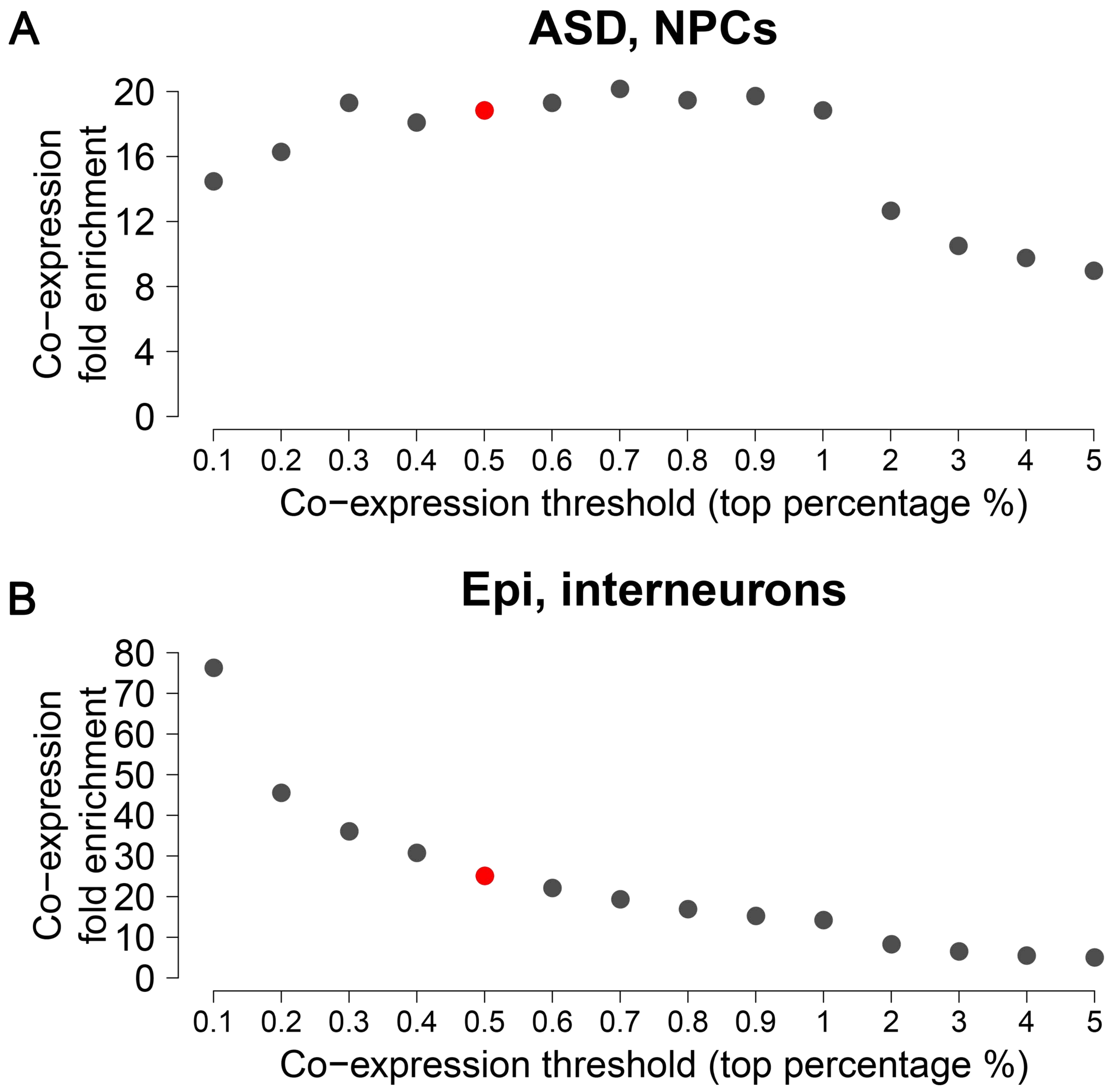


**Supplementary Figure S6.** Co-expression enrichment analysis of dnLoF-ASD genes in NPCs and dnMis-Epi genes in interneurons at varying co-expression thresholds. (*A,B*) Co-expression fold enrichment of dnLoF-ASD genes in NPCs (*A*) and dnMis-Epi genes in interneurons (*B*) at varying co-expression thresholds between top 0.1% and top 5%. Circle size is proportional to co-expression fold enrichment score.

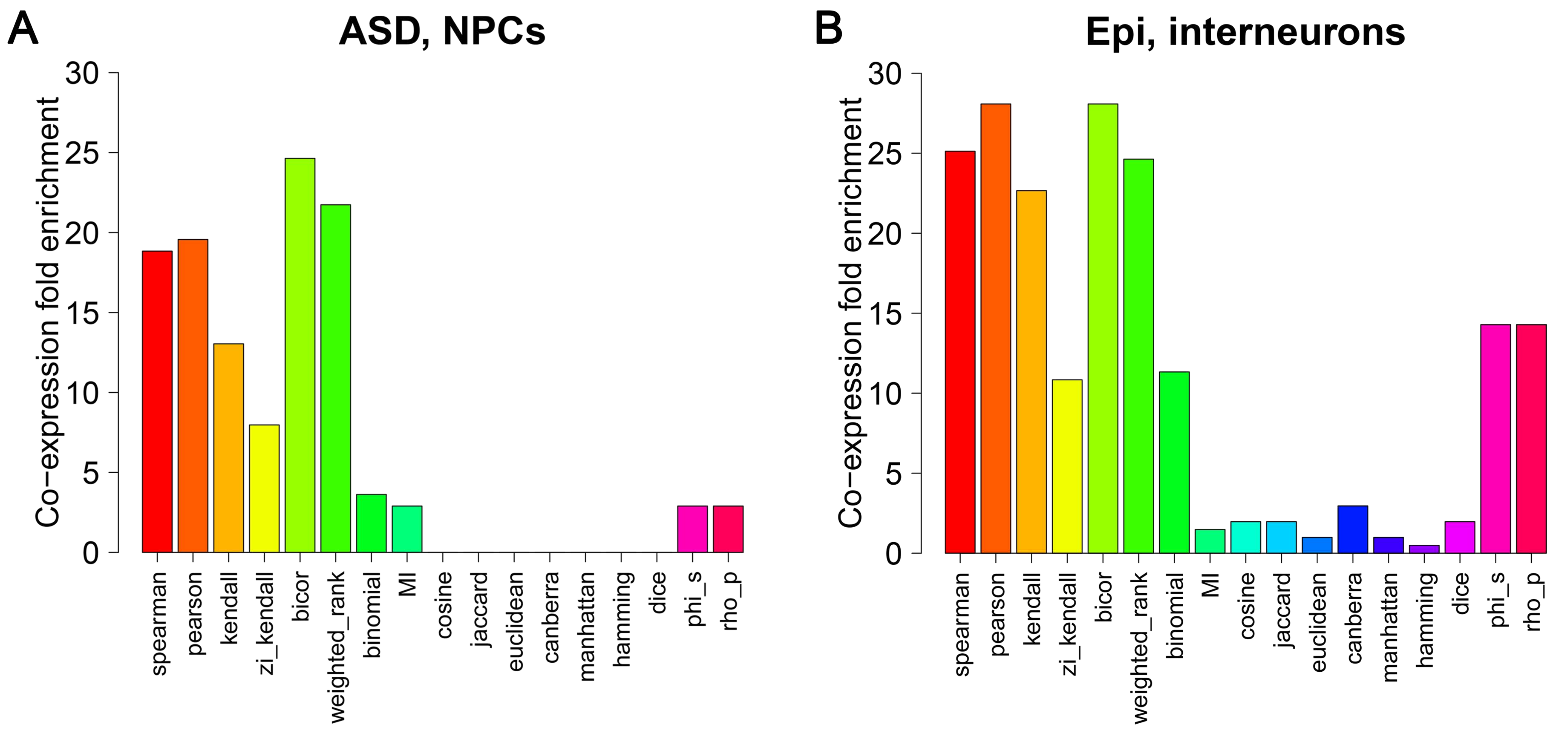


**Supplementary Figure S7.** Co-expression enrichment analysis of dnLoF-ASD genes in NPCs and dnMis-Epi genes in interneurons using different measures of association. (*A,B*) Co-expression fold enrichment of dnLoF-ASD genes in NPCs (*A*) and dnMis-Epi genes in interneurons (*B*) using 17 measures of association implemented in the 'dismay' R package.

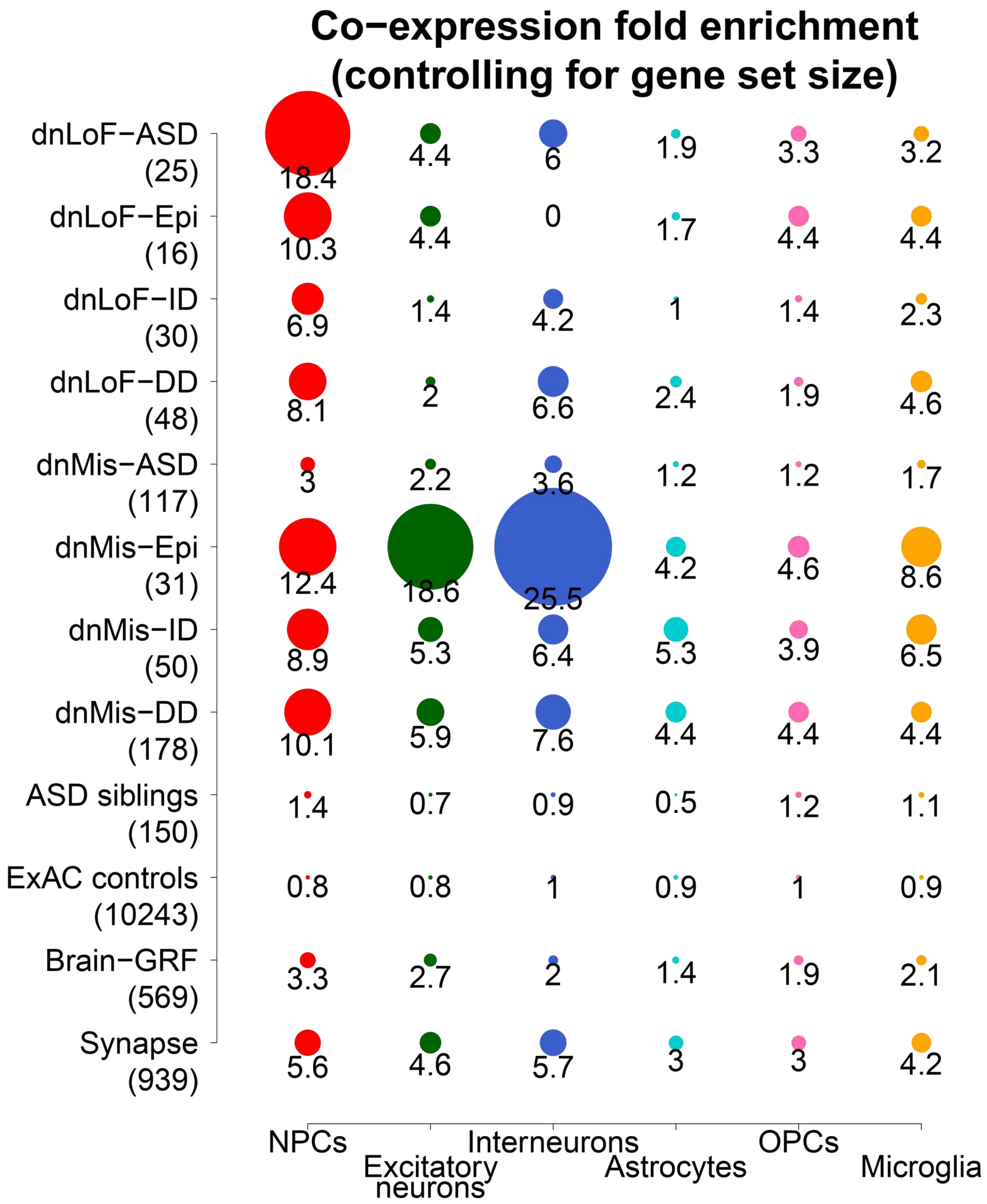


**Supplementary Figure S8.** Co-expression enrichment analysis of twelve gene sets in six major cell types of human prefrontal cortex by controlling for gene set size. Co-expression fold enrichment of eight NDD gene sets and four control gene sets in six major cortical cell types by controlling for gene set size. For all the twelve gene sets in each cell type, co-expression enrichment score is computed using randomly chosen twelve gene sets with the same gene set size (the minimum gene set size of the original twelve gene sets in the cell type) from the original twelve gene sets 1000 times. Gene set size is shown in parentheses. Circle size is proportional to co-expression fold enrichment score.

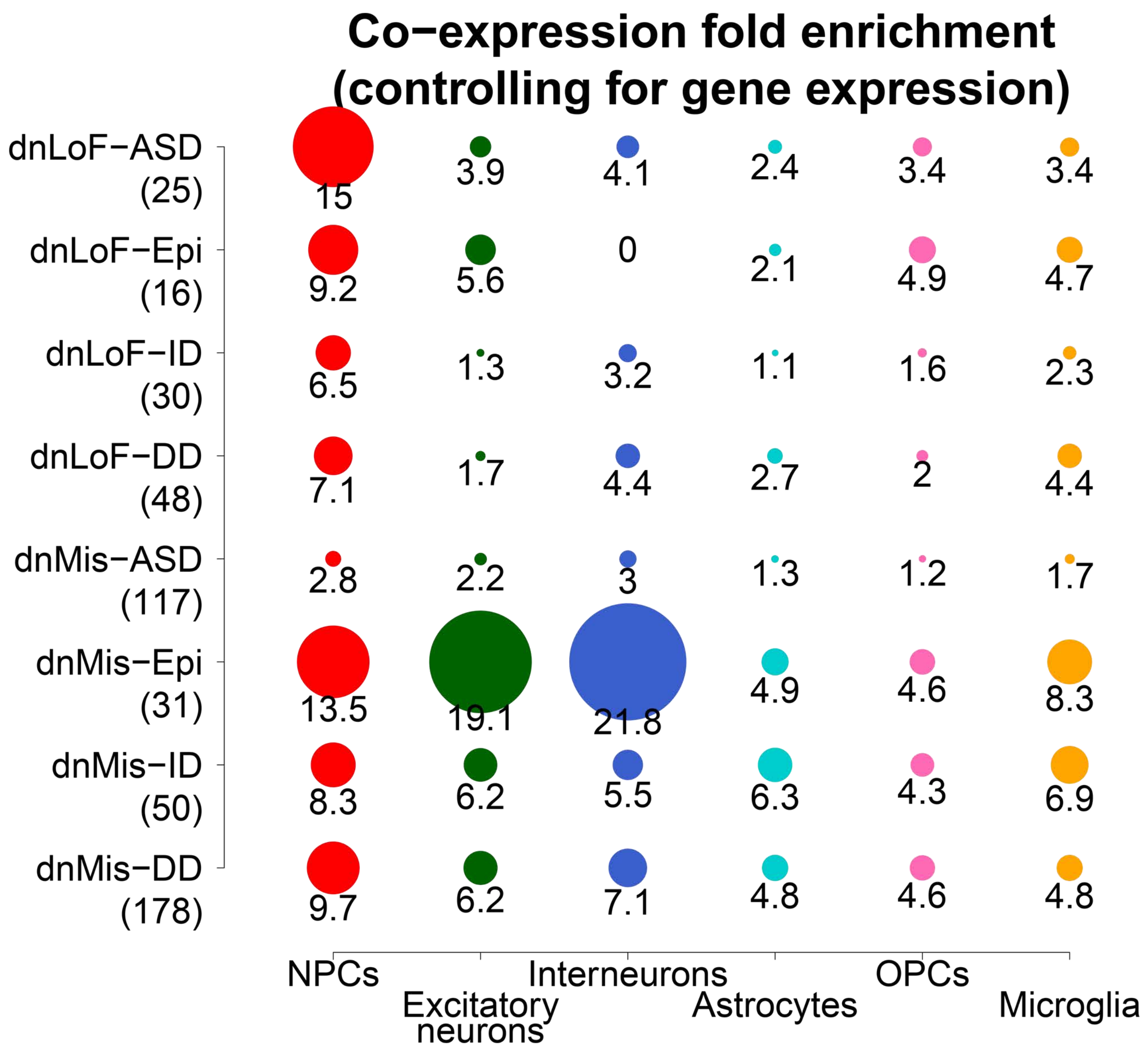


**Supplementary Figure S9.** Co-expression enrichment analysis of eight NDD gene sets in six major cell types of human prefrontal cortex by controlling for expression level dependence. Co-expression fold enrichment of eight NDD gene sets in six major cortical cell types by controlling for expression level dependence. For each gene set in each cell type, co-expression enrichment score is computed using 1000 randomly chosen gene sets with similar expression levels in the cell type as the background gene set. Gene set size is shown in parentheses. Circle size is proportional to co-expression fold enrichment score.

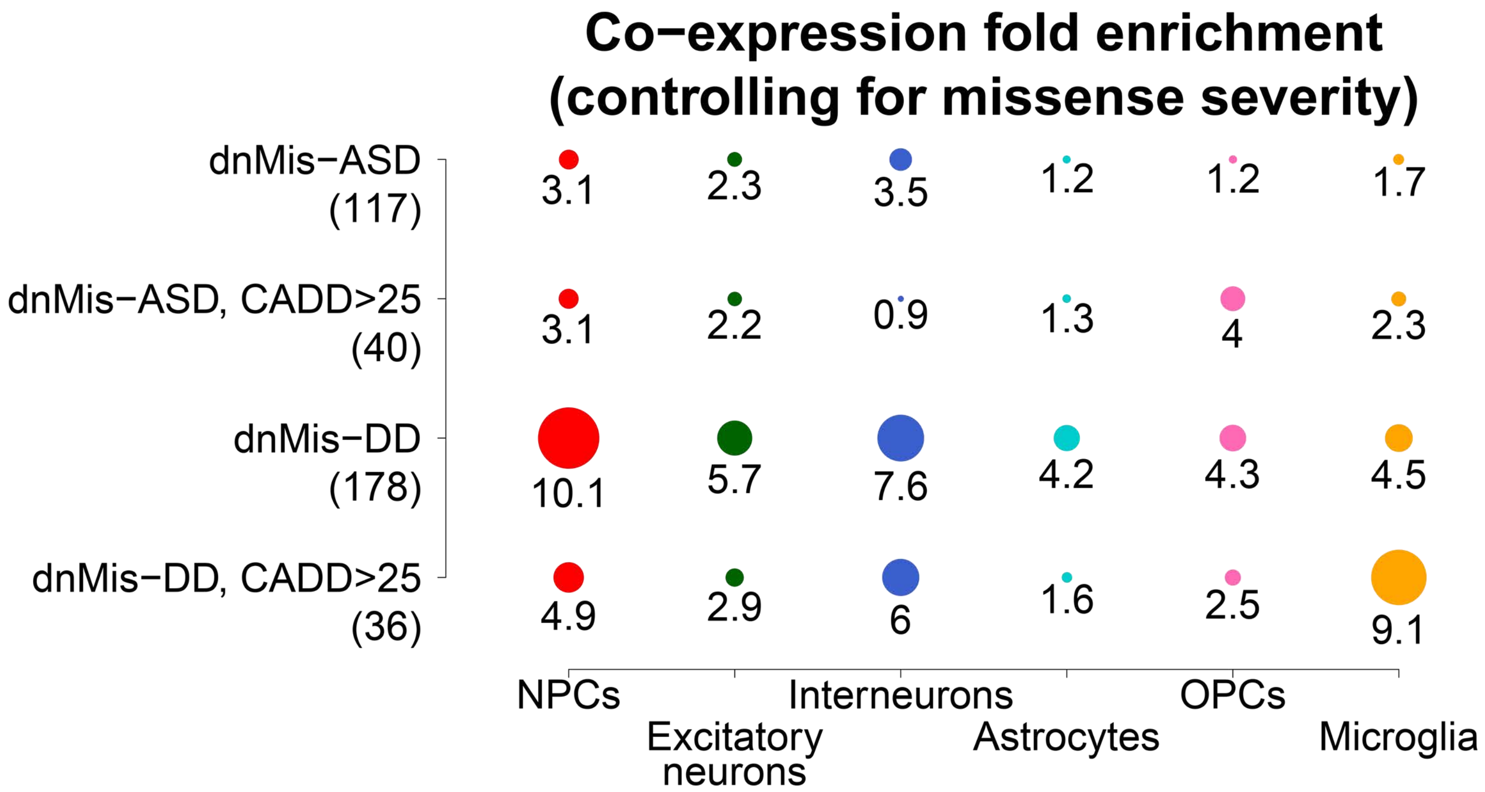


**Supplementary Figure S10.** Co-expression enrichment analysis of ASD and DD gene sets with dnMis mutations in six major cell types of human prefrontal cortex by controlling for missense severity. Co-expression fold enrichment of ASD and DD gene sets with dnMis mutations and those with severe dnMis mutations (CADD score>25) in six major cortical cell types. Gene set size is shown in parentheses. Circle size is proportional to co-expression fold enrichment score.

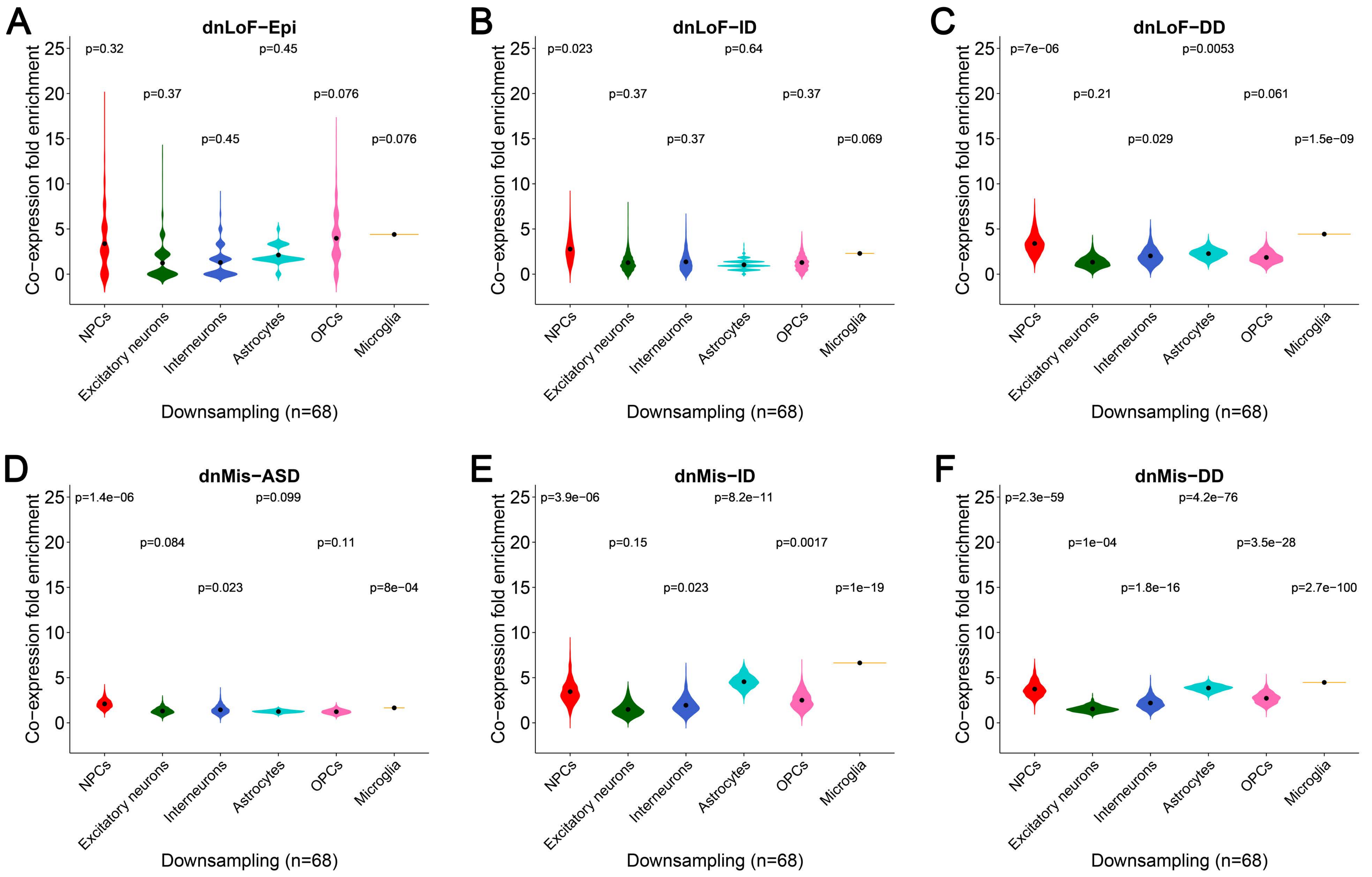


**Supplementary Figure S11.** Co-expression enrichment analysis of the other six NDD gene sets in six major cell types of human prefrontal cortex by downsampling. (*A-F*) Co-expression fold enrichment of dnLoF-Epi (*A*), dnLoF-ID (*B*), dnLoF-DD (*C*), dnMis-ASD (*D*), dnMis-ID (*E*), and dnMis-DD genes (*F*) in six major cortical cell types by downsampling the same number of cells for each cell type. Violin plot shows the mean value (point). The statistical significance P value measures whether the mean co-expression fold enrichment score of the corresponding gene set is higher than that of the background genes by the one-sided Fisher's exact test.

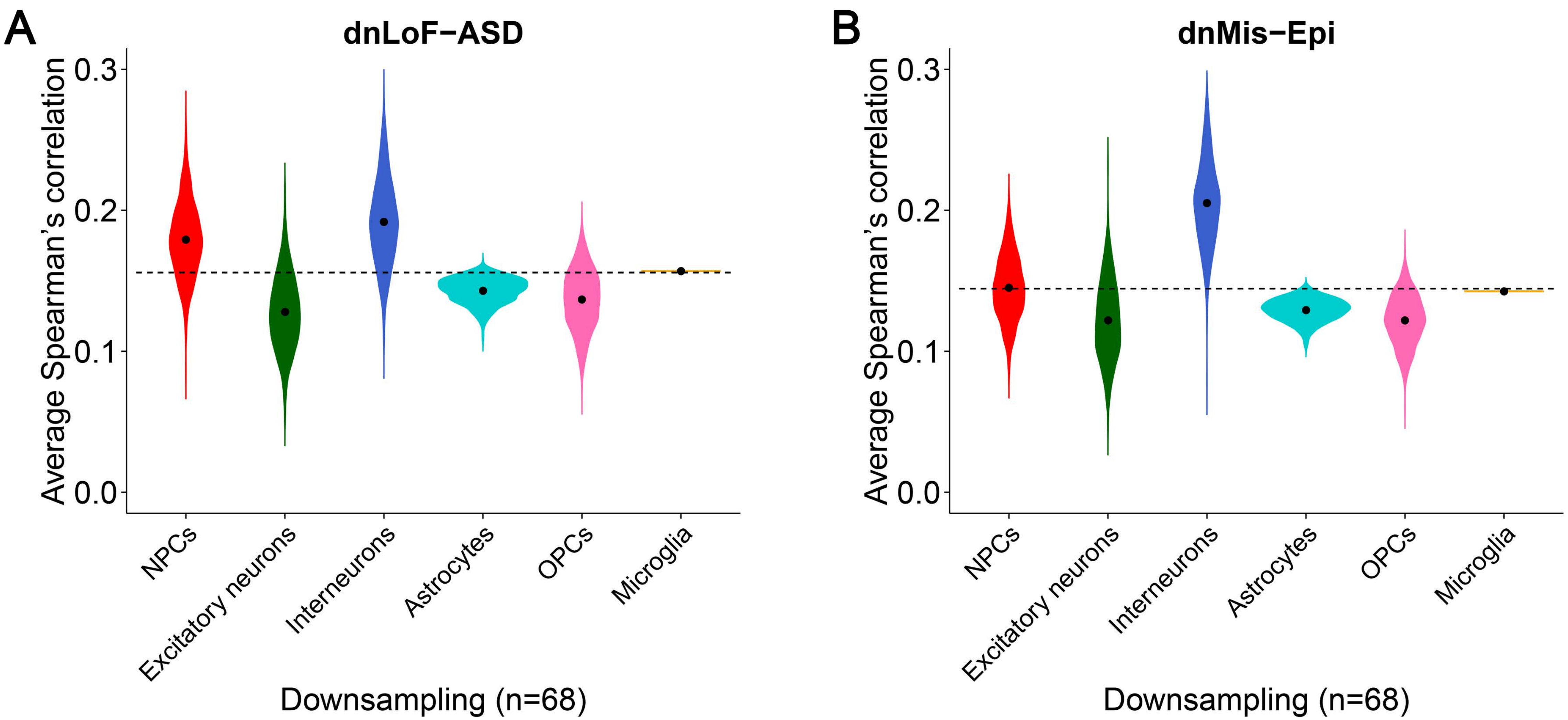


**Supplementary Figure S12.** Spearman's correlation analysis of dnLoF-ASD and dnMis-Epi genes in six major cell types of human prefrontal cortex by downsampling. (*A,B*) Average Spearman's correlation of dnLoF-ASD (*A*) and dnMis-Epi genes (*B*) in six major cortical cell types by downsampling the same number of cells for each cell type. Violin plot shows the mean value (point). The dashed horizontal line indicates the mean of six mean values of 1000 average Spearman's correlation coefficients of an NDD gene set for six major cell types by downsampling.

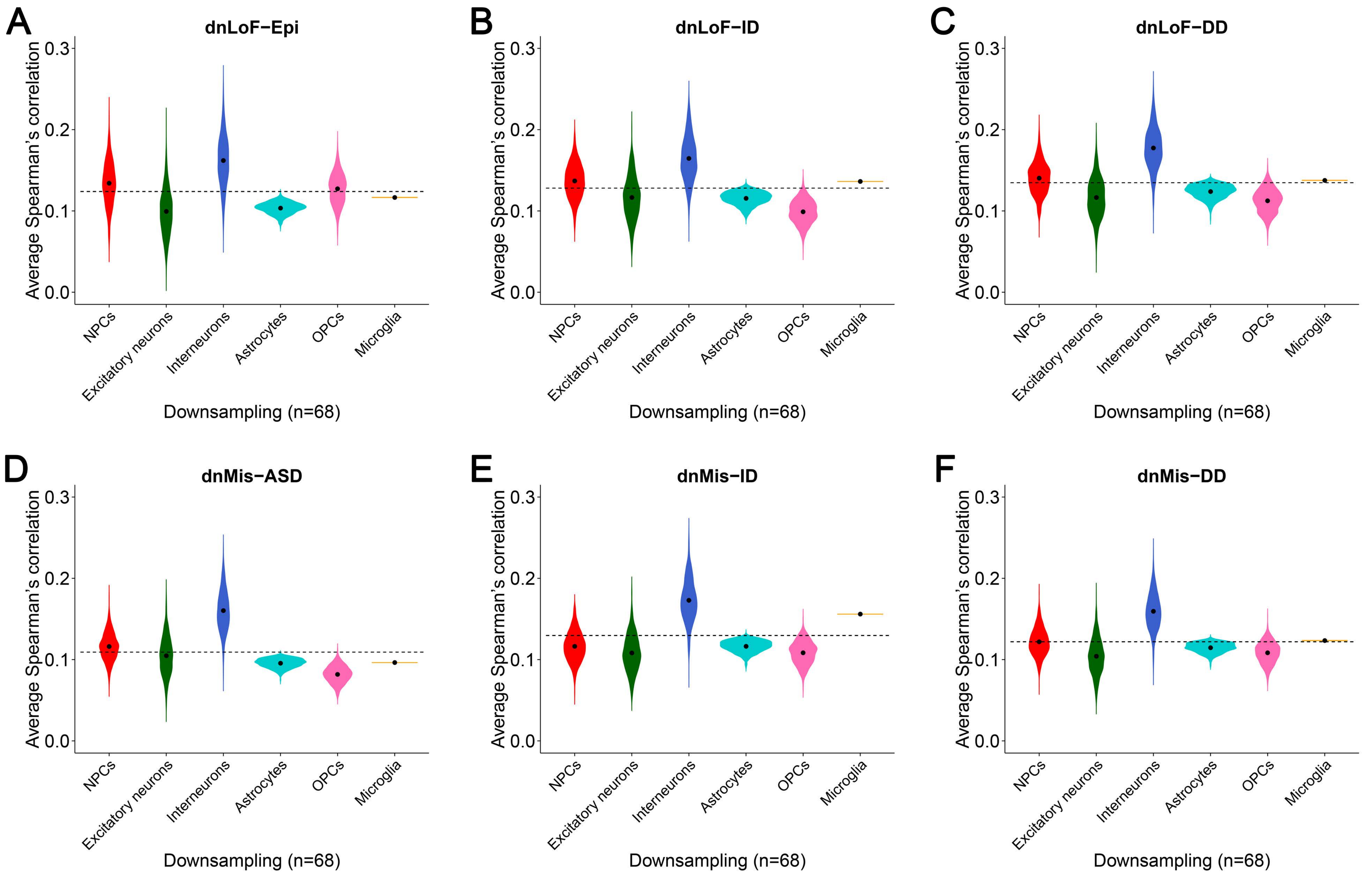


**Supplementary Figure S13.** Spearman's correlation analysis of the other six NDD gene sets in six major cell types of human prefrontal cortex by downsampling. (*A-F*) Average Spearman's correlation of dnLoF-Epi (*A*), dnLoF-ID (*B*), dnLoF-DD (*C*), dnMis-ASD (*D*), dnMis-ID (*E*), and dnMis-DD genes (*F*) in six major cortical cell types by downsampling the same number of cells for each cell type. Violin plot shows the mean value (point). The dashed horizontal line indicates the mean of six mean values of 1000 average Spearman's correlation coefficients of an NDD gene set for six major cell types by downsampling.

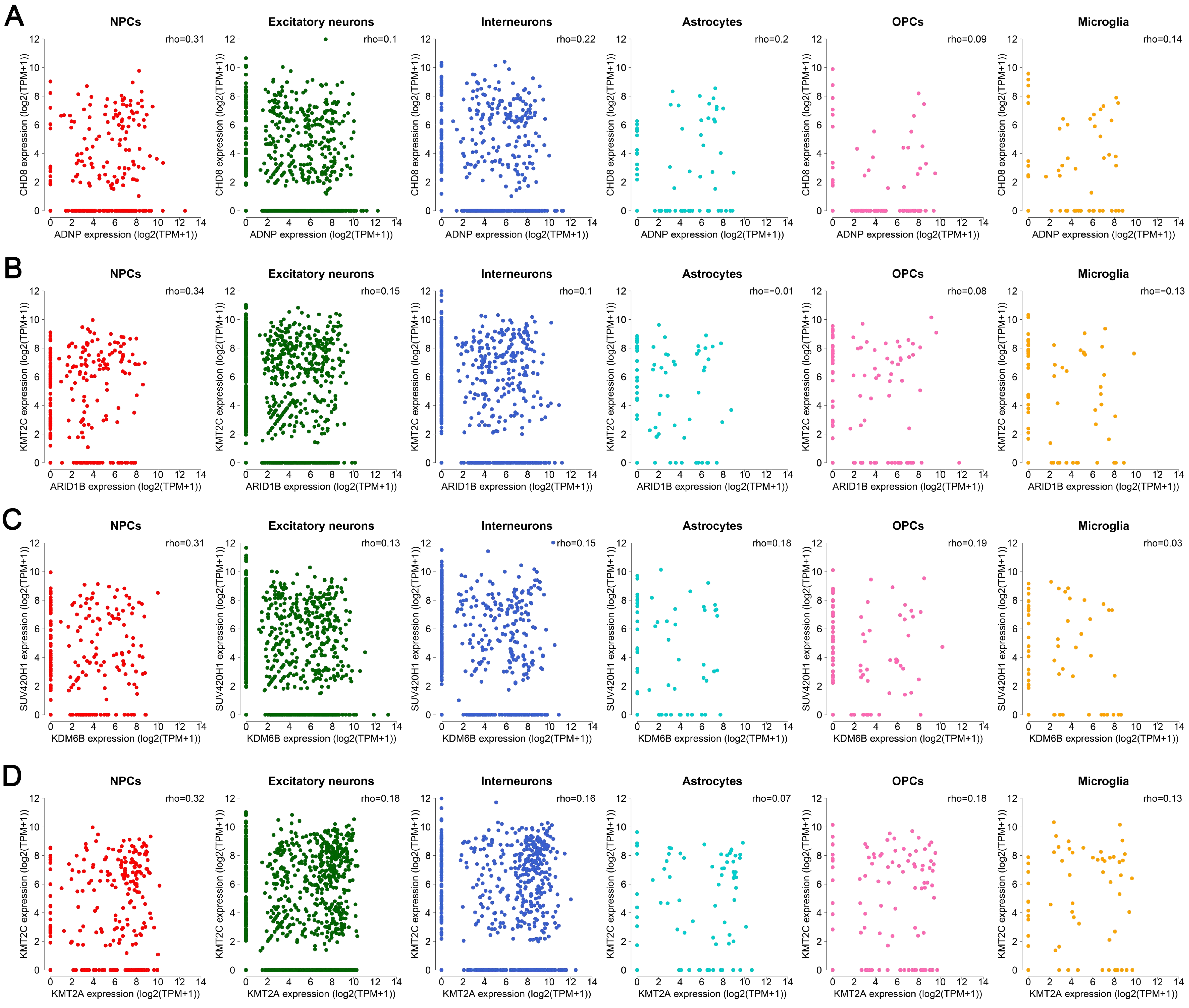


**Supplementary Figure S14.** Examples of dnLoF-ASD gene pairs that show higher co-expression in NPCs than in the other five major cell types. (*A-D*) Scatter plots show the expression levels of one dnLoF-ASD gene versus the expression levels of another dnLoF-ASD gene in six major cell types. The mean values of 1000 Spearman's correlation coefficients by downsampling are shown.

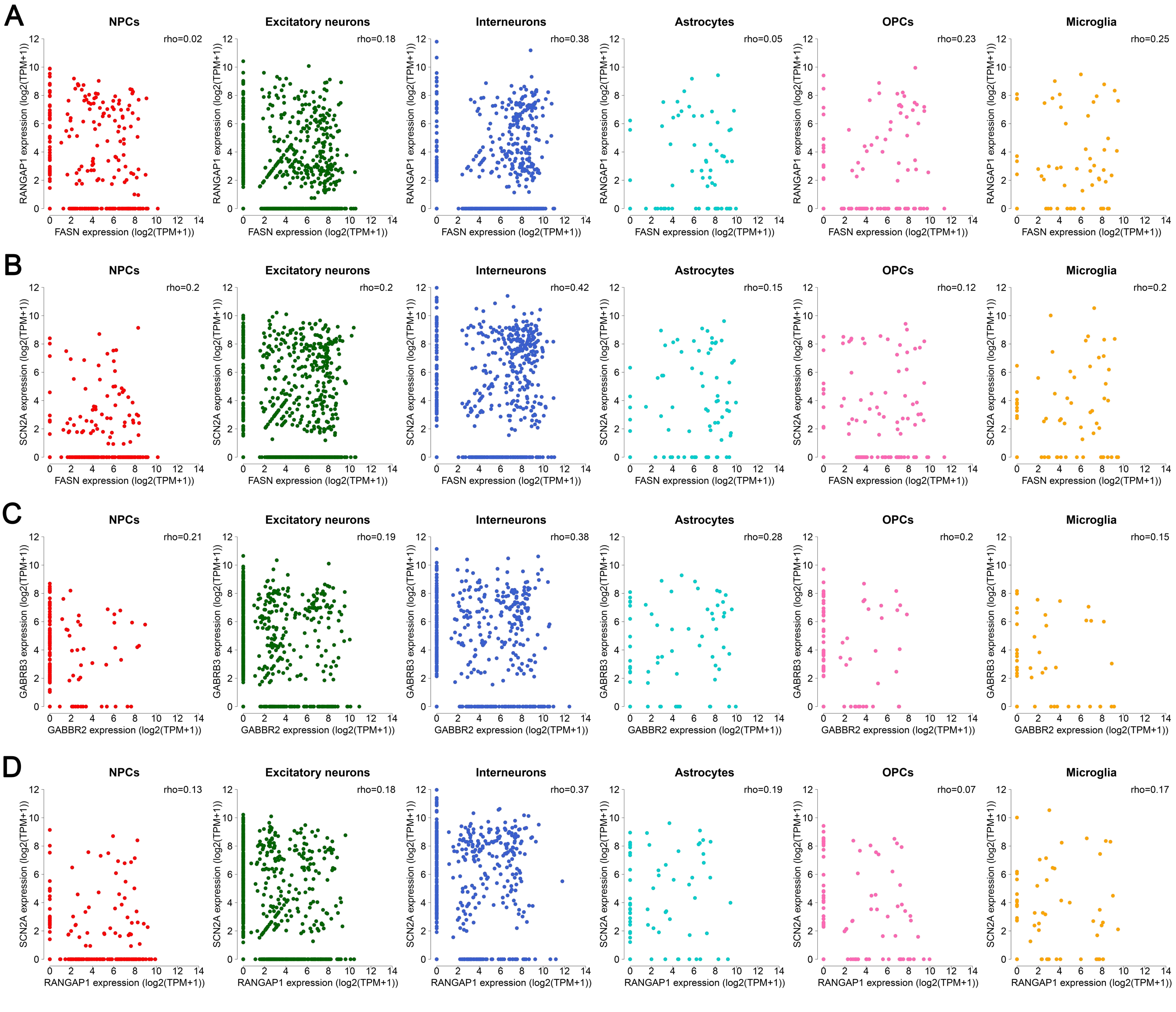


**Supplementary Figure S15.** Examples of dnMis-Epi gene pairs that show higher co-expression in interneurons than in the other five major cell types. (*A-D*) Scatter plots show the expression levels of one dnMis-Epi gene versus the expression levels of another dnMis-Epi gene in six major cell types. The mean values of 1000 Spearman's correlation coefficients by downsampling are shown.

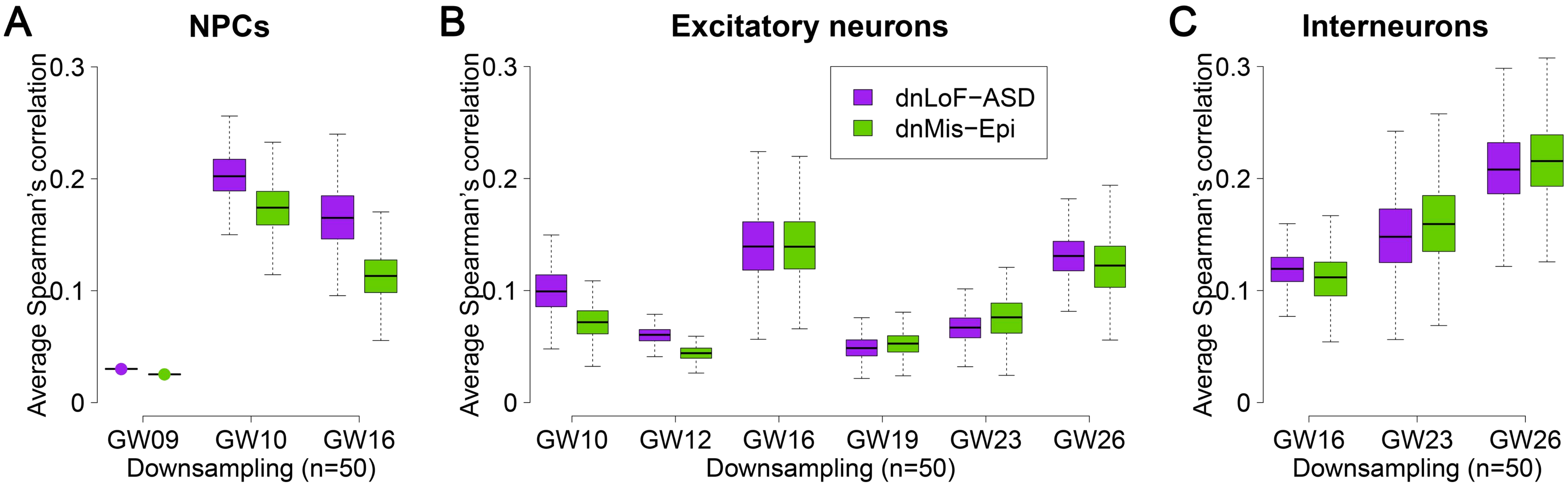


**Supplementary Figure S16.** Spearman's correlation analysis of dnLoF-ASD and dnMis-Epi genes during NPC and neuron development by downsampling. (*A-C*) Average Spearman's correlation of dnLoF-ASD and dnMis-Epi genes at specific stages of NPCs (*A*), excitatory neurons (*B*), and interneurons (*C*) by downsampling the same number of cells for each cell stage.

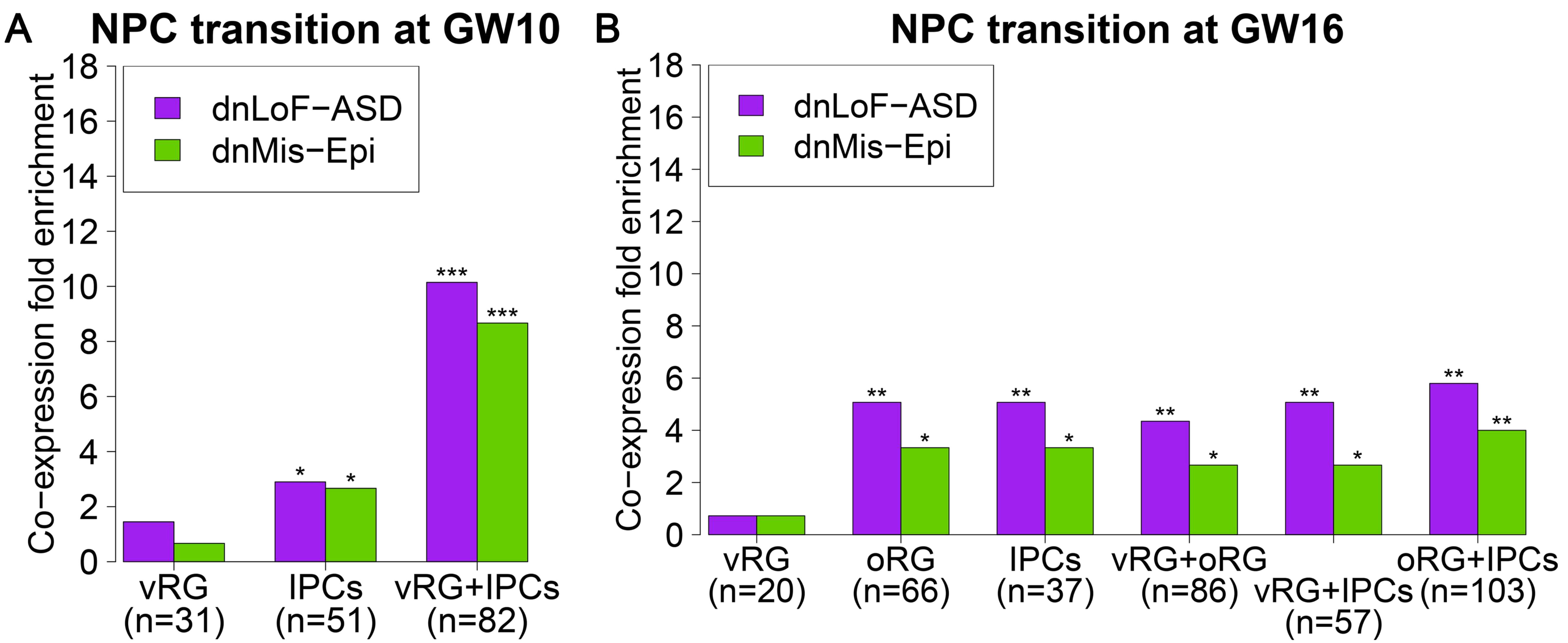


**Supplementary Figure S17.** Co-expression enrichment analysis of dnLoF-ASD and dnMis-Epi genes during NPC transitions. (*A*) Co-expression fold enrichment of dnLoF-ASD and dnMis-Epi genes in vRG cells, IPCs, and the transition at GW10 using the original number of cells which is shown in parentheses. (*B*) Co-expression fold enrichment of dnLoF-ASD and dnMis-Epi genes in vRG cells, oRG cells, IPCs, and their transitions at GW16 using the original number of cells which is shown in parentheses. In (*A,B*), asterisks above bar indicate -log10(P) value that measures statistical significance whether the corresponding gene set has a higher co-expression fold enrichment score than the background genes by the one-sided Fisher's exact test (* 1≤ -log10(P) <2, ** 2≤ -log10(P) <5, *** 5≤ -log10(P) <10, **** 10≤ -log10(P)).

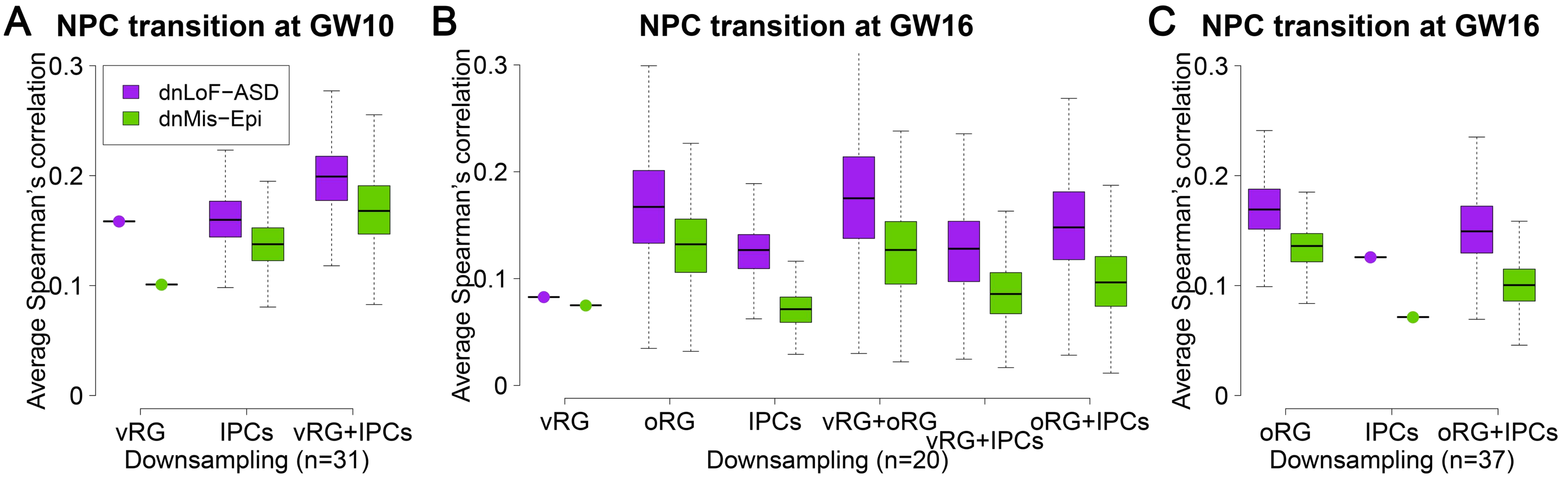


**Supplementary Figure S18.** Spearman's correlation analysis of dnLoF-ASD and dnMis-Epi genes during NPC transitions by downsampling. (*A*) Average Spearman's correlation of dnLoF-ASD and dnMis-Epi genes in vRG cells, IPCs, and the transition at GW10 by downsampling the same number of cells for each condition. (*B*) Average Spearman's correlation of dnLoF-ASD and dnMis-Epi genes in vRG cells, oRG cells, IPCs, and their transitions at GW16 by downsampling the same number of cells for each condition. (*C*) Average Spearman's correlation of dnLoF-ASD and dnMis-Epi genes in oRG cells, IPCs, and the transition at GW16 by downsampling the same number of cells for each condition.

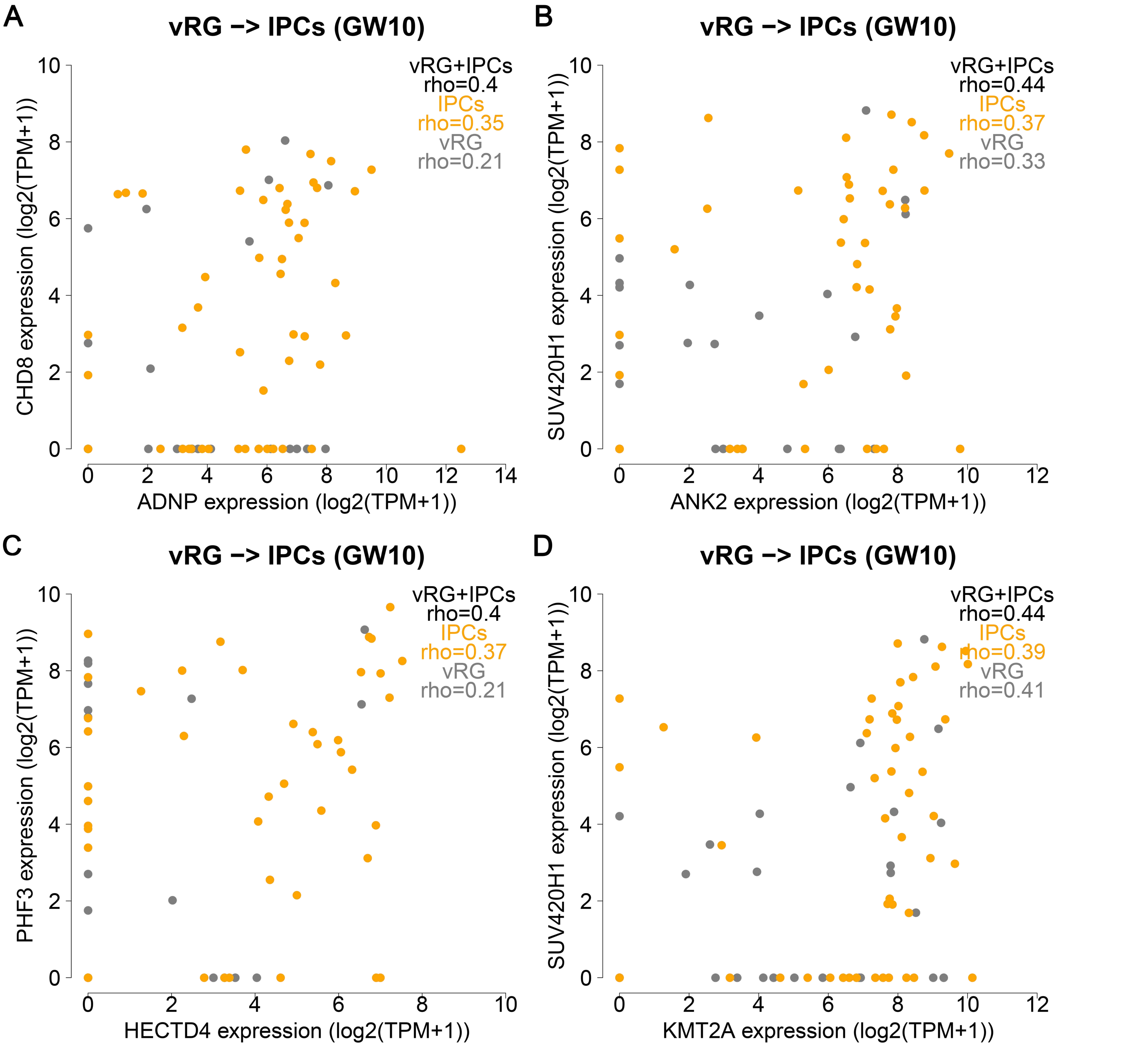


**Supplementary Figure S19.** Examples of dnLoF-ASD gene pairs that show increased co-expression during the vRG-to-IPC transition at GW10 (vRG cells and IPCs combined) compared with vRG cells or IPCs alone. (*A-D*) Scatter plot shows the expression levels of one dnLoF-ASD gene versus the expression levels of another dnLoF-ASD gene. The mean value of 1000 Spearman's correlation coefficients by downsampling is shown.

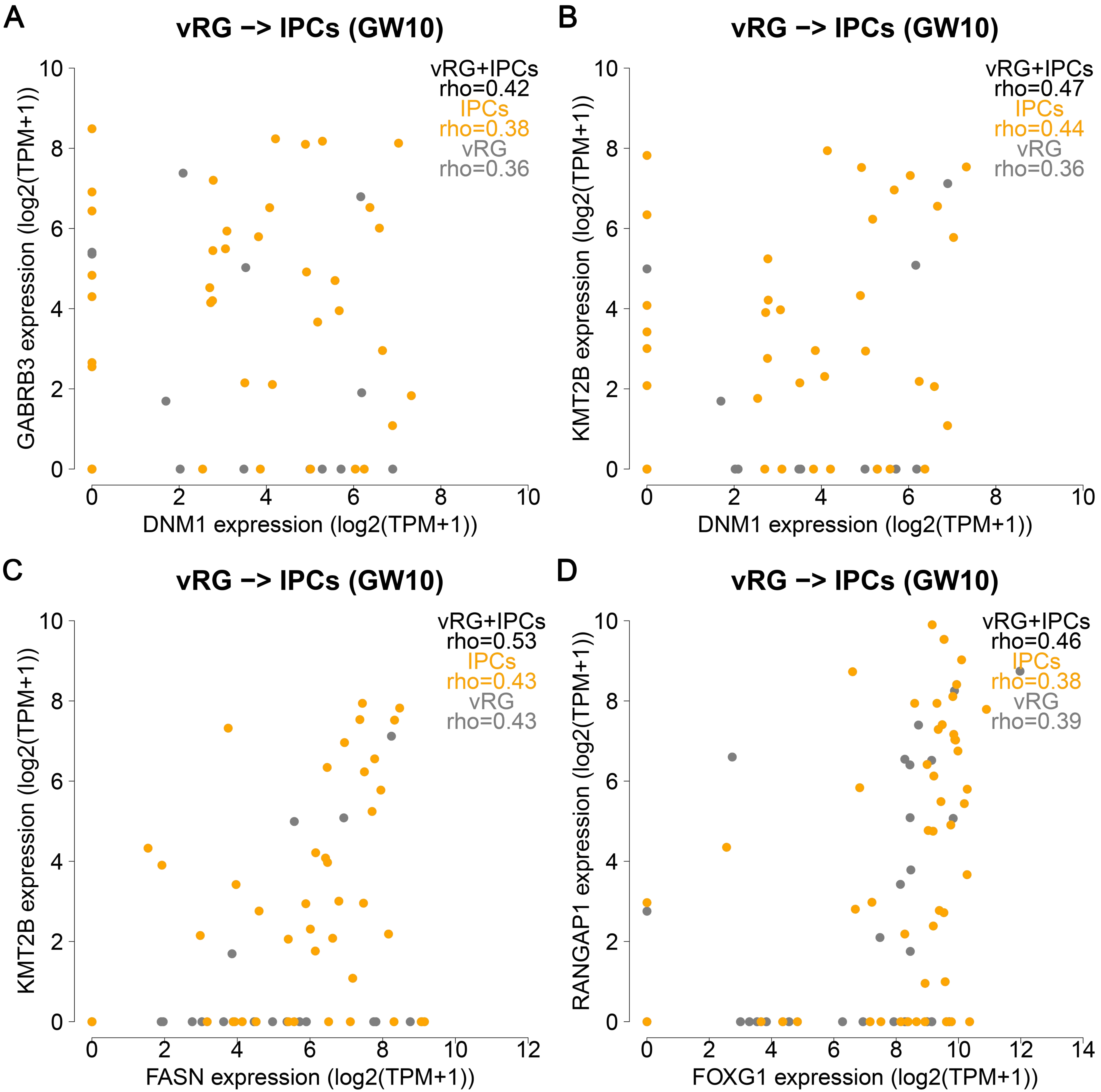


**Supplementary Figure S20.** Examples of dnMis-Epi gene pairs that show increased co-expression during the vRG-to-IPC transition at GW10 (vRG cells and IPCs combined) compared with vRG cells or IPCs alone. (*A-D*) Scatter plot shows the expression levels of one dnMis-Epi gene versus the expression levels of another dnMis-Epi gene. The mean value of 1000 Spearman's correlation coefficients by downsampling is shown.

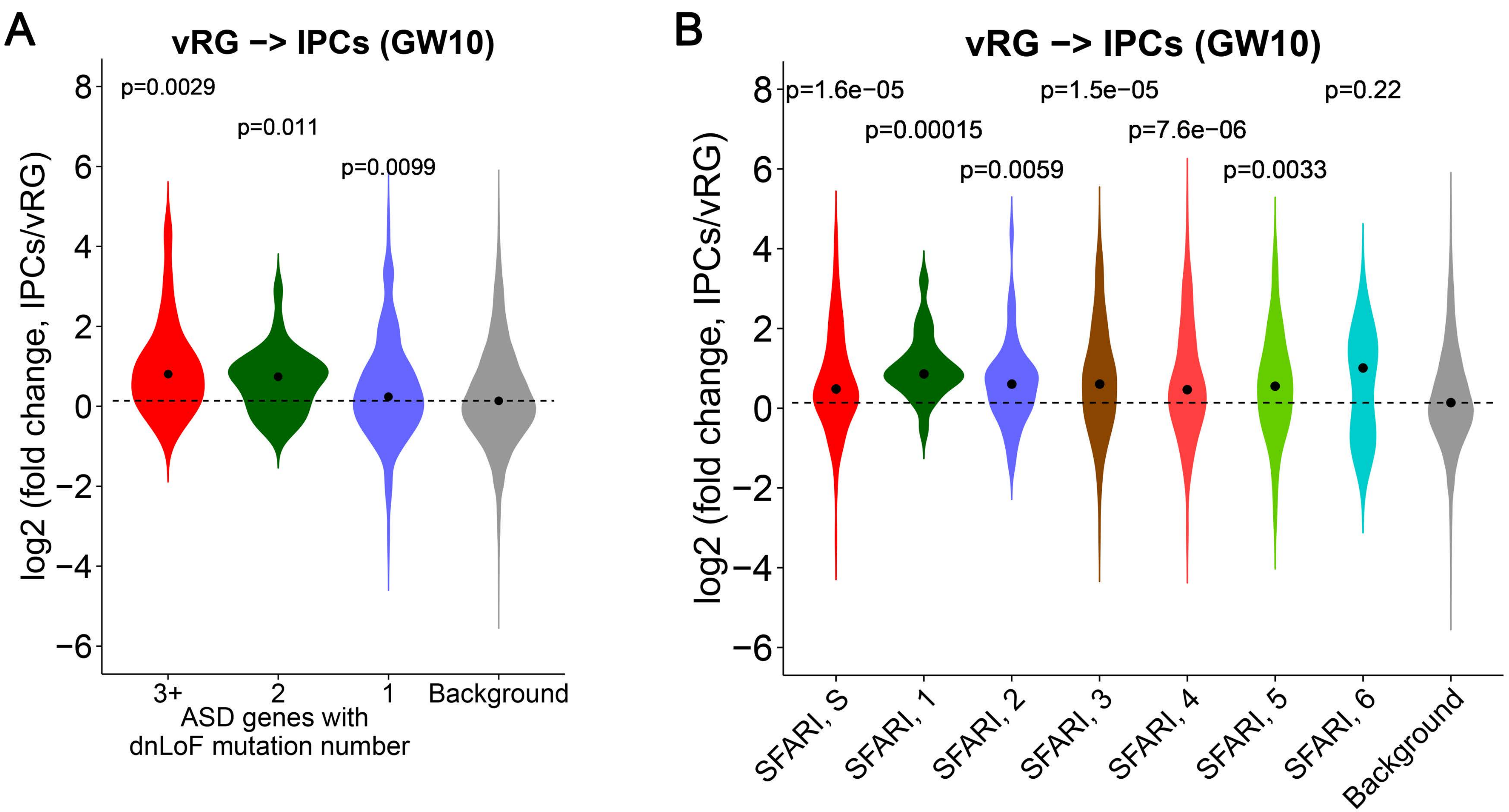


**Supplementary Figure S21.** Expression change of diverse ASD gene sets during the transition from vRG cells to IPCs at GW10. (*A*) The expression change of ASD genes with ≥3, 2 and 1 dnLoF mutations during the transition. The dashed horizontal line indicates median log2(fold change) value of the background genes. The statistical significance P values measure whether different types of ASD genes with dnLoF mutations have higher log2(fold change) values than the background genes by the one-sided Wilcoxon rank sum test. (*B*) The expression change of different types of SFARI ASD genes during the transition. The dashed horizontal line indicates the median log2(fold change) value of the background genes. The statistical significance P values measure whether different types of SFARI ASD genes have higher log2(fold change) values than the background genes by the one-sided Wilcoxon rank sum test.

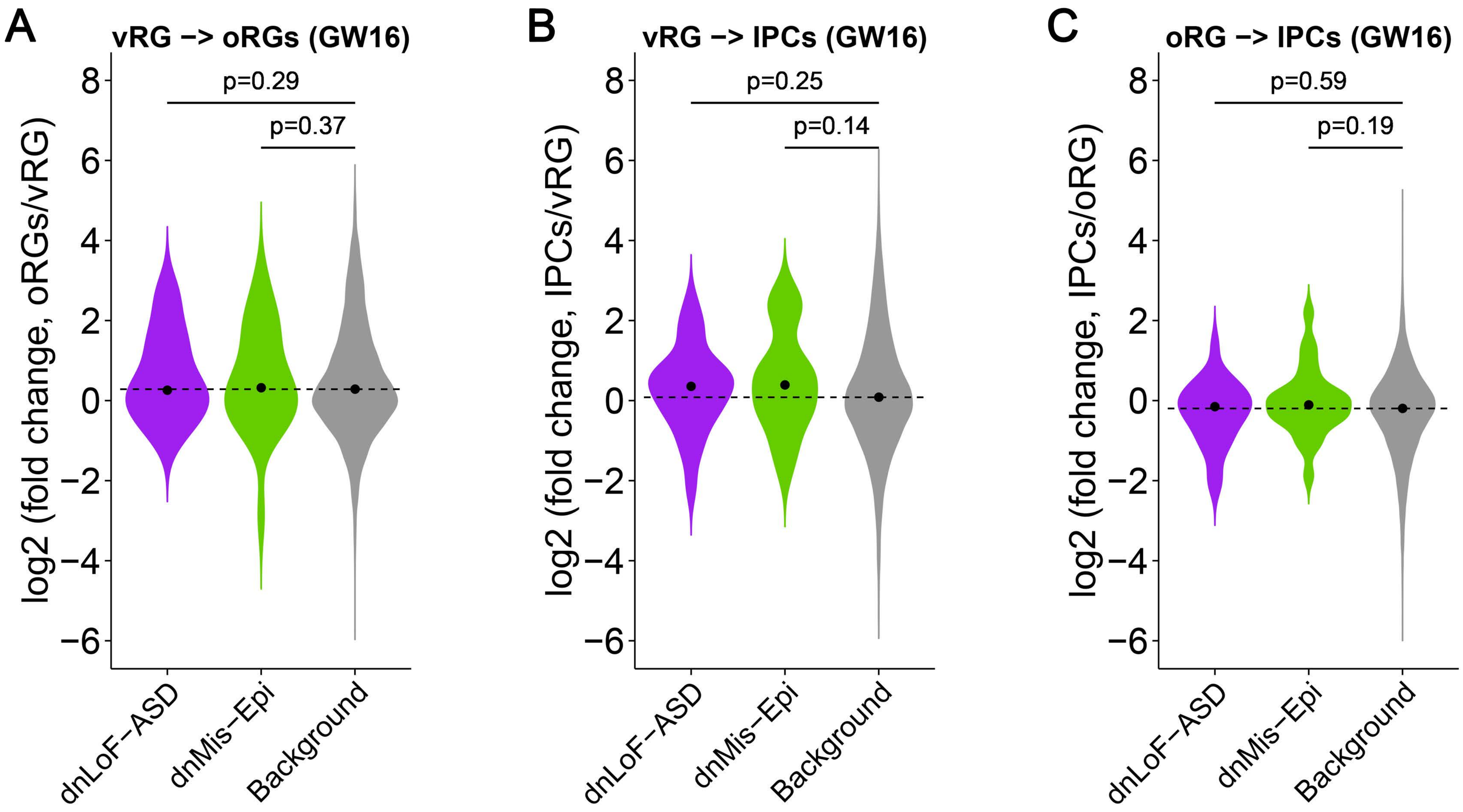


**Supplementary Figure S22.** Expression change of dnLoF-ASD and dnMis-Epi genes during the cell type transitions within NPCs at GW16. (*A*-*C*) The expression change of dnLoF-ASD and dnMis-Epi genes during the transitions at GW16 from vRG cells to oRG cells (*A*), vRG cells to IPCs (*B*), and oRG cells to IPCs (*C*). The dashed horizontal line indicates the median log2(fold change) value of the background genes. The statistical significance P values measure whether dnLoF-ASD and dnMis-Epi genes have higher log2(fold change) values than the background genes during the transitions by the one-sided Wilcoxon rank sum test.

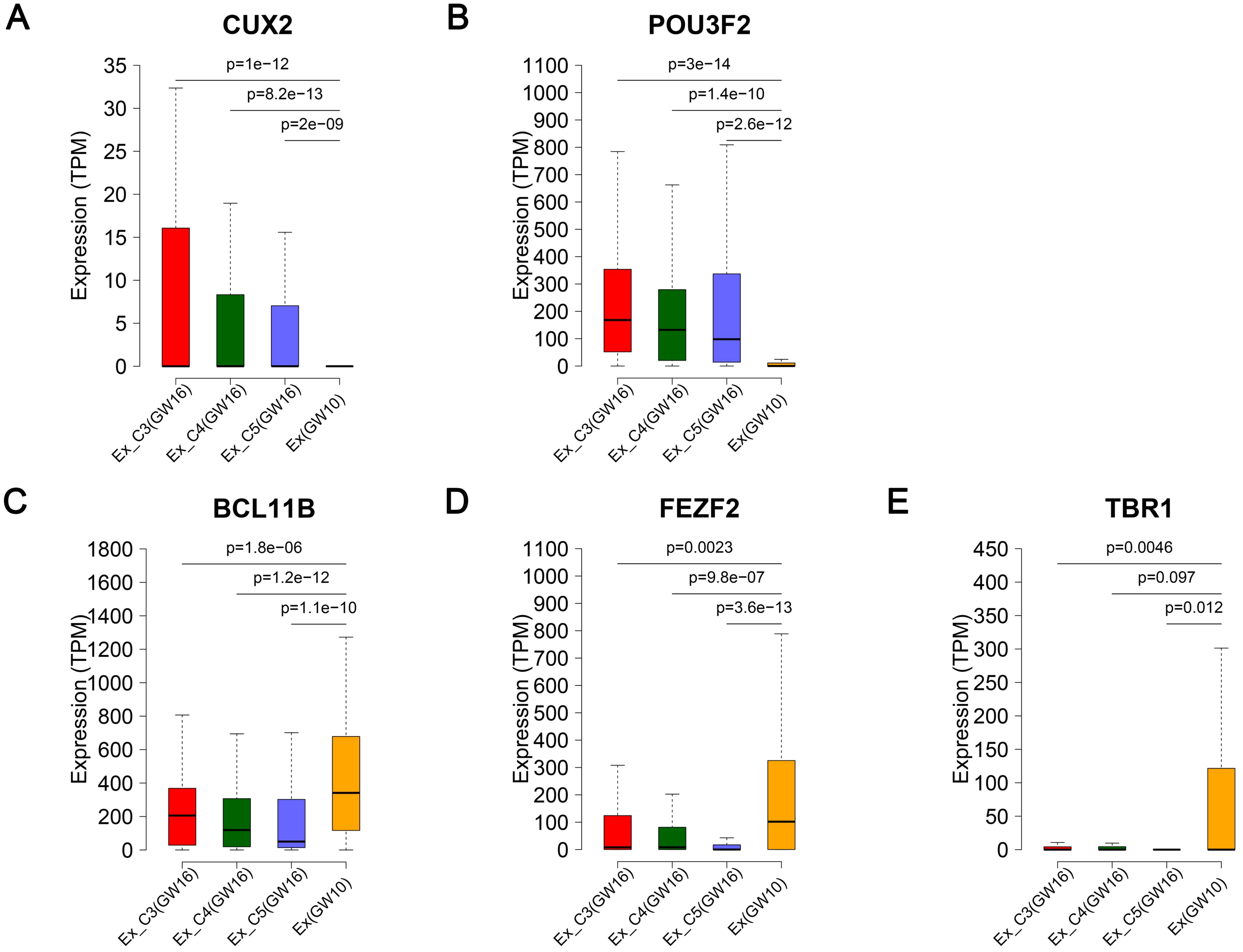


**Supplementary Figure S23.** Expression difference of layer marker genes in subclusters of excitatory neurons (Ex) at GW16 compared with excitatory neurons at GW10. (*A-E*) The expression difference of upper-layer neuron markers *CUX2* (*A*), and *POU3F2* (*B*), and deep-layer neuron markers *BCL11B* (*C*), *FEZF2* (*D*), and *TBR1* (*E*) in the three subclusters of excitatory neurons at GW16 compared with excitatory neurons at GW10. The statistical significance P values measure expression difference of layer marker genes between GW16 excitatory neuron subclusters and GW10 excitatory neurons using DESeq2.

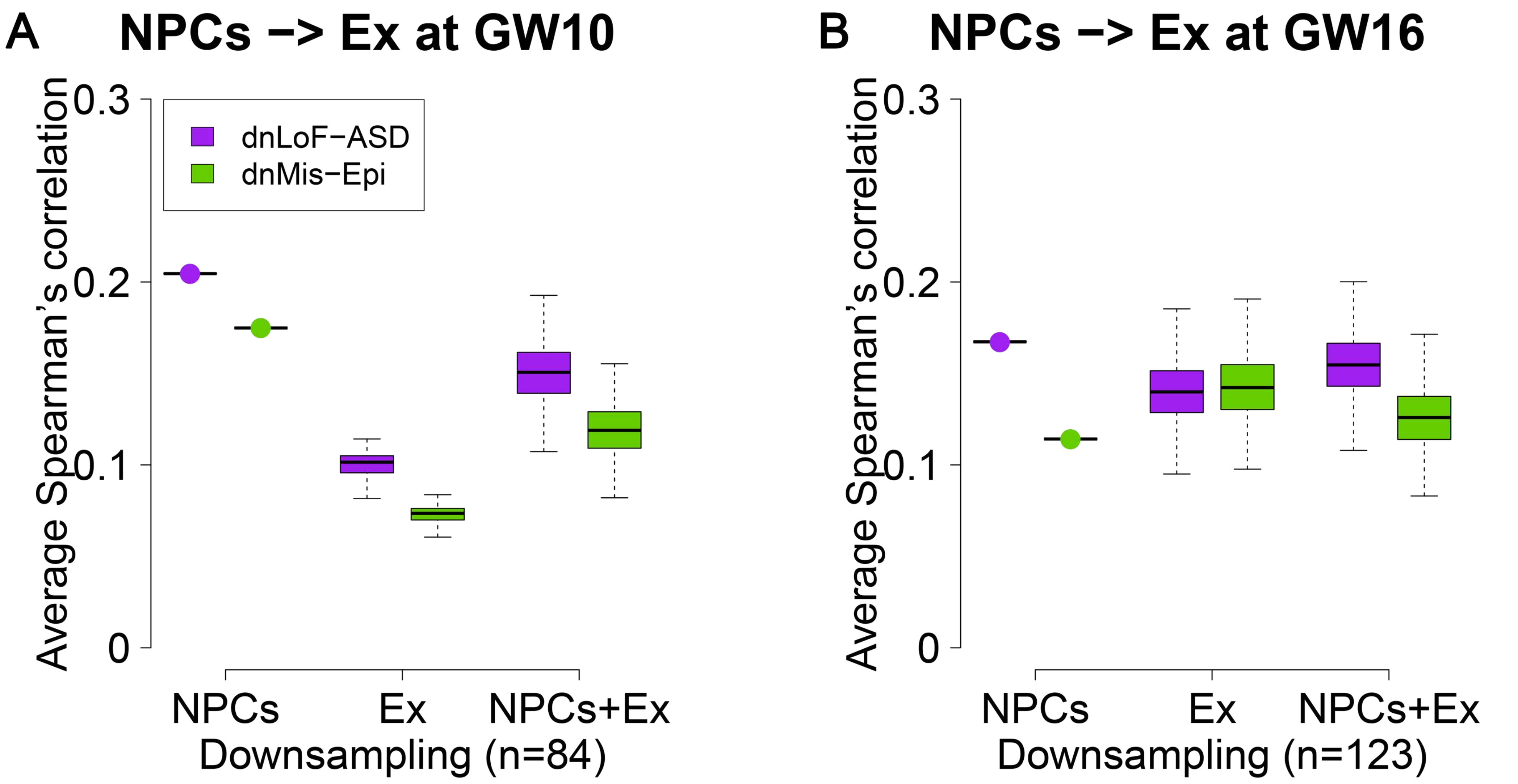


**Supplementary Figure S24.** Spearman's correlation analysis of dnLoF-ASD and dnMis-Epi genes during differentiation from NPCs to excitatory neurons (Ex) by downsampling. (*A,B*) Average Spearman's correlation of dnLoF-ASD and dnMis-Epi genes in NPCs, excitatory neurons, and the differentiation at GW10 (*A*) and GW16 (*B*) by downsampling the same number of cells for each condition.

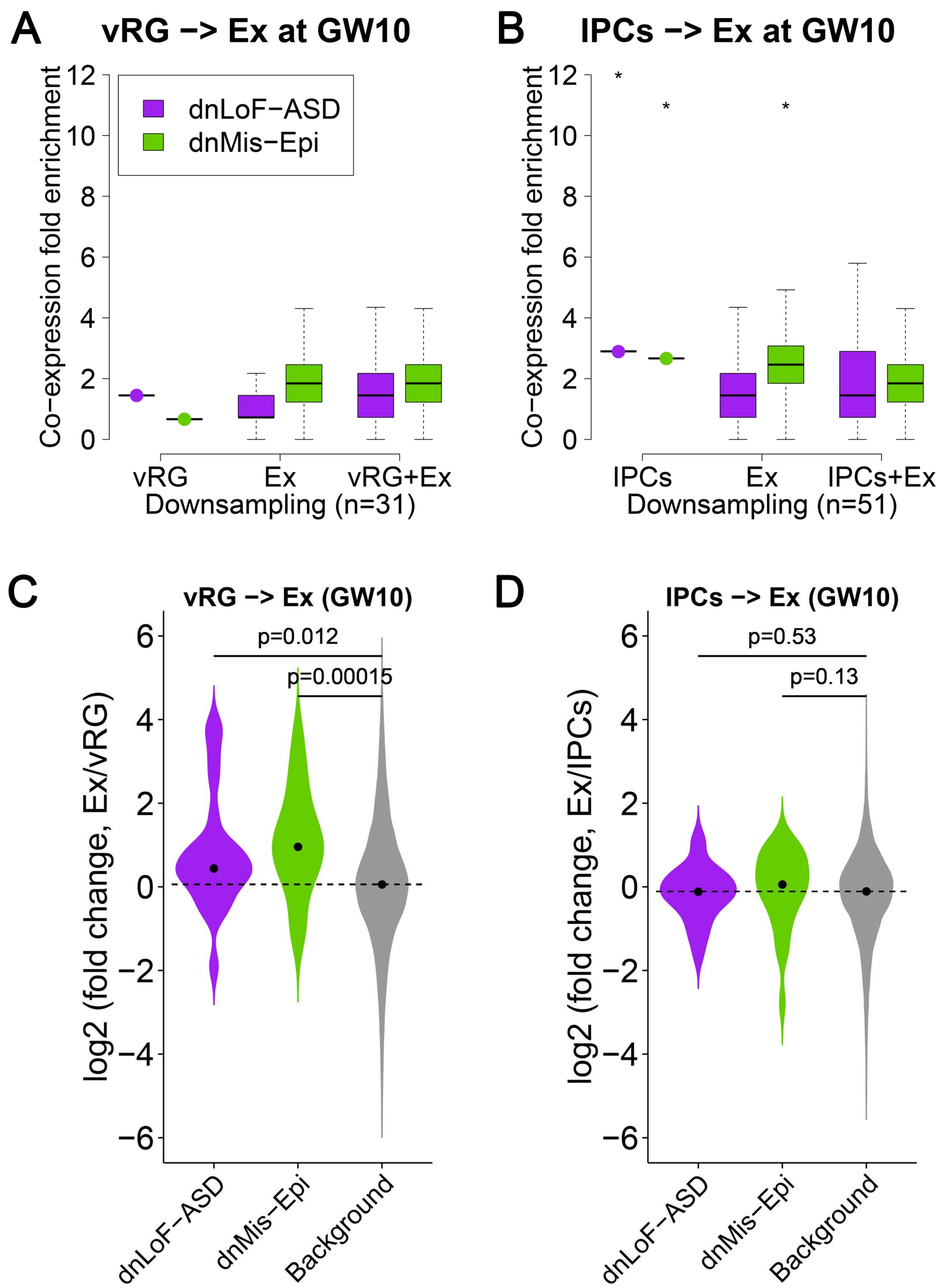


**Supplementary Figure S25.** Co-expression enrichment analysis of dnLoF-ASD and dnMis-Epi genes during the differentiation from vRG cells and IPCs to excitatory neurons (Ex) at GW10. (*A,B*) Co-expression fold enrichment of dnLoF-ASD and dnMis-Epi genes in vRG cells, excitatory neurons, and the differentiation at GW10 (*A*) and in IPCs, excitatory neurons, and the differentiation at GW10 (*B*) by downsampling the same number of cells for each condition. Asterisks above boxplot indicate -log10(P) value that measures statistical significance whether the mean co-expression fold enrichment score of the corresponding gene set is higher than that of the background genes by the one-sided Fisher's exact test (* 1≤ -log10(P) <2, ** 2≤ -log10(P) <5, *** 5≤ -log10(P) <10, **** 10≤ -log10(P)). (*C,D*) The expression change of dnLoF-ASD and dnMis-Epi genes during the differentiation at GW10 from vRG cells to excitatory neurons (*C*) and IPCs to excitatory neurons (*D*). The dashed horizontal line indicates the median log2(fold change) value of the background genes. The statistical significance P values measure whether dnLoF-ASD and dnMis-Epi genes have higher log2(fold change) values than the background genes during the differentiation by the one-sided Wilcoxon rank sum test.

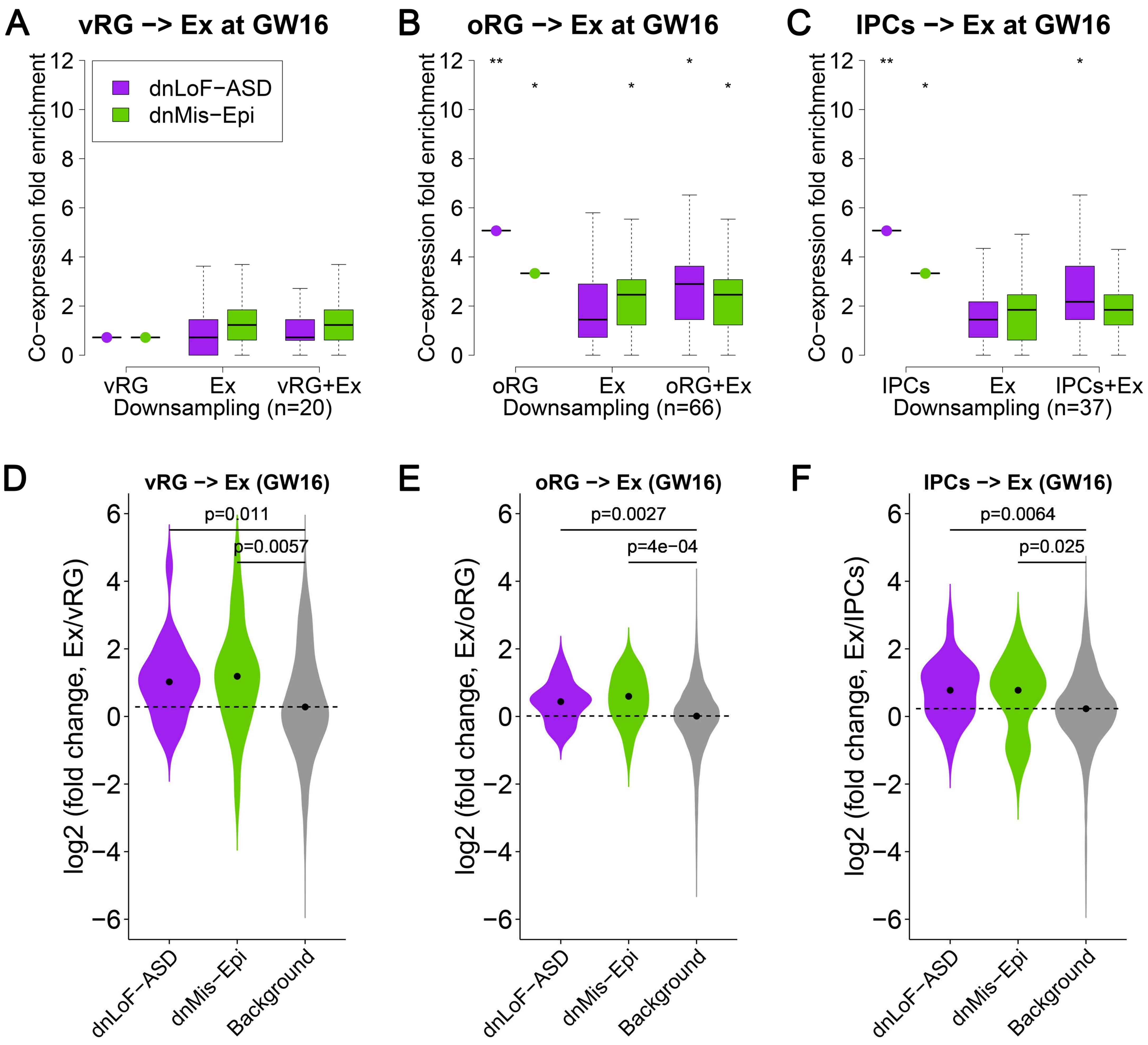


**Supplementary Figure S26.** Co-expression enrichment analysis of dnLoF-ASD and dnMis-Epi genes during the differentiation from vRG cells, oRG cells, and IPCs to excitatory neurons (Ex) at GW16. (*A-C*) Co-expression fold enrichment of dnLoF-ASD and dnMis-Epi genes in vRG cells, excitatory neurons, and the differentiation at GW16 (*A*), in oRG cells, excitatory neurons, and the differentiation at GW16 (*B*), and in IPCs, excitatory neurons, and the differentiation at GW16 (*C*) by downsampling the same number of cells for each condition. Asterisks above boxplot indicate -log10(P) value that measures statistical significance whether the mean co-expression fold enrichment score of the corresponding gene set is higher than that of the background genes by the one-sided Fisher's exact test (* 1≤ -log10(P) <2, ** 2≤ -log10(P) <5, *** 5≤ -log10(P) <10, **** 10≤ -log10(P)). (*D-F*) The expression change of dnLoF-ASD and dnMis-Epi genes during the differentiation at GW16 from vRG cells to excitatory neurons (*D*), oRG cells to excitatory neurons (*E*), and IPCs to excitatory neurons (*F*). The dashed horizontal line indicates the median log2(fold change) value of the background genes. The statistical significance P values measure whether dnLoF-ASD and dnMis-Epi genes have higher log2(fold change) values than the background genes during the differentiation by the one-sided Wilcoxon rank sum test.

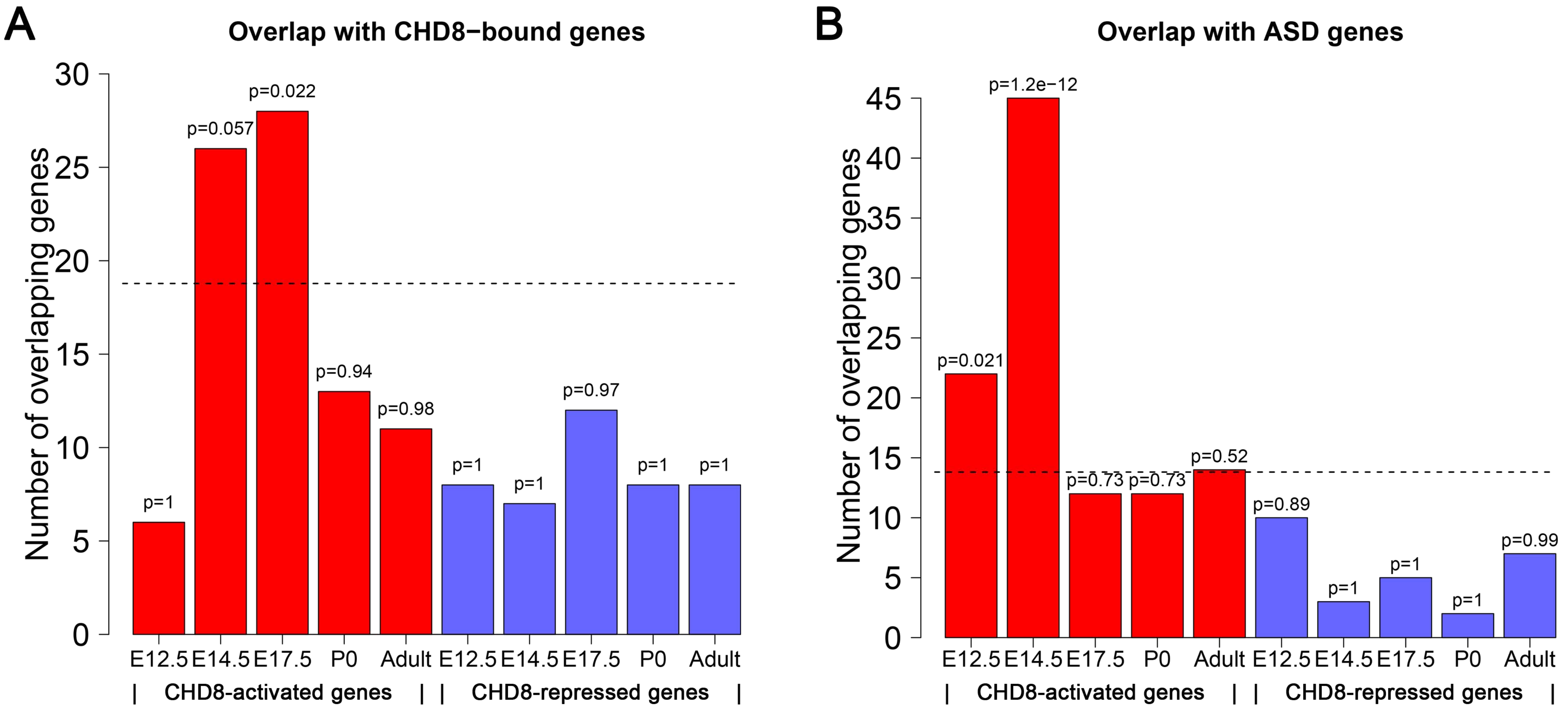


**Supplementary Figure S27.** *CHD8* target gene selection. (*A*) Overlap between *CHD8*-activated/-repressed genes and *CHD8*-bound genes. The dashed horizontal line indicates the expected number of *CHD8*-bound genes among 300 random genes from the background genes. The statistical significance P values measure whether *CHD8*-activated and -repressed genes are enriched with *CHD8*-bound genes by the one-sided Fisher's exact test. (*B*) Overlap between *CHD8*-activated/-repressed genes and ASD genes with at least one dnLoF mutations. The dashed horizontal line indicates the expected number of ASD genes among 300 random genes from the background genes. The statistical significance P values measure whether *CHD8*-activated and -repressed genes are enriched with ASD genes by the one-sided Fisher's exact test.
